# Epithelial QKI Protects Against Emphysema by Maintaining Mitochondrial Integrity

**DOI:** 10.64898/2026.07.09.737340

**Authors:** Kiyoshi Uemasu, Kazuya Tanimura, Atsushi Miyamoto, Koichi Hasegawa, Zachary Lane, Reika Nyunoya, Haruka Uemasu, Brett A. Kaufman, Corrine R. Kliment, Divay Chandra, Frank Sciurba, Charles Dela Cruz, Prithu Sundd, Jonathan K. Alder, Jian Hu, Toru Nyunoya

## Abstract

Single-cell transcriptomic profiling of chronic obstructive pulmonary disease (COPD) lungs identified QKI, an RNA-binding protein, as a candidate emphysema-associated gene, but its epithelial role in COPD pathobiology remains unclear. We show that QKI expression is reduced in human COPD lungs and that alveolar type 2 epithelial (AT2) cell QKI protein levels correlate strongly with spirometric indices and diffusing capacity (DL_CO_). Lung epithelium-specific QKI knockout mice (QKI^Δ/Δ^) developed spontaneous airspace enlargement with emphysema-like mechanics, and QKI-deficient AT2 cells showed impaired spheroid colony formation and increased apoptosis. Integrated transcriptomic and proteomic analyses of primary AT2 cells revealed a selective reduction in functional mitochondrial (respiratory-chain and metabolic) protein abundance despite relatively preserved transcript levels, consistent with mitochondrial transcriptome-proteome discordance. QKI loss increased mtDNA abundance and TOMM20 staining but decreased ATP5A, indicating accumulation of structurally increased but functionally dysfunctional mitochondria. In human epithelial cells, CRISPR-mediated QKI deficiency reduced oxidative respiration, increased glycolytic reliance, elevated mitochondrial ROS and membrane potential, and increased apoptosis; these phenotypes were partially rescued by QKI re-expression. These findings identify epithelial QKI as a regulator of mitochondrial integrity and stress tolerance in COPD.

## Introduction

Chronic obstructive pulmonary disease (COPD) is characterized by an irreversible airflow limitation and is the fourth leading cause of death worldwide (1). COPD, often caused by cigarette smoking, is a heterogeneous disorder with distinct phenotypes, each likely involving unique biological mechanisms (2). Among these, emphysema is defined by the destructive enlargement of the airspaces and is a key phenotype contributing to high morbidity and mortality in patients (3–6).

Although multiple mechanisms, including cell apoptosis, senescence, oxidative stress, and protease-antiprotease imbalance, have been implicated, the epithelial-intrinsic factors that make some individuals more susceptible to CS-induced emphysema remain unclear (7–11). Our single cell RNA sequencing analysis of COPD lungs identified *QKI*, a K homology domain-containing RNA-binding protein (RBP), as a COPD candidate gene (12) among 127 emphysema-associated genes with altered expression in lung parenchyma based on emphysema severity (13).

We previously reported that QKI protein expression is significantly reduced in smokers with COPD compared with smokers without the disease (12). QKI regulates key biological functions, including cell proliferation, differentiation, and anti-inflammatory responses, by controlling RNA metabolism at both co- and post-transcriptional levels – affecting mRNA splicing, transport, stability, and micro/circular RNA formation (14–18). However, the role of QKI in maintaining alveolar epithelial integrity remains poorly understood.

Of its three major isoforms (QKI-5, QKI-6, and QKI-7), QKI-5 is the predominant form in lung cells (15, 19). Functionally, QKI is essential in the central nervous system, where it regulates oligodendrocyte differentiation and myelination (20, 21), and in the vascular system, where it maintains endothelial stability, angiogenesis, and cardiac myofibrogenesis (22–24). QKI also acts as a tumor suppressor in lung cancer (25). Despite its broad expression across lung cell types – including bronchial and alveolar epithelial cells, fibroblasts, endothelial cells, and alveolar macrophages (12) – the role of QKI in COPD remains poorly understood.

This study aimed to investigate the relationship between QKI expression in human lung tissues and clinical parameters using two complementary approaches. First, we analyzed QKI gene expression across a broad cohort, including individuals with and without COPD, as well as smokers and non-smokers. Second, we examined alveolar type II (AT2) cell-specific QKI protein expression in surgically resected lung tissues obtained from smokers with or without COPD. Building on these human data, we then determined whether QKI loss in lung epithelial cells (LECs) alone is sufficient to drive spontaneous emphysematous changes in an experimental mouse model. By integrating clinical and translational data, we aimed to determine whether QKI expression correlates with disease severity in human COPD and whether QKI is essential for maintaining alveolar integrity *in vivo*.

## Results

### QKI Expression is Reduced in COPD Lungs and Associated with Impaired Lung Function

To investigate the role of QKI in COPD, we examined QKI gene expression in whole lung tissue samples from individuals with and without COPD (patient characteristics are shown in Table E2). QKI expression was significantly decreased in COPD, with severe to very severe COPD (GOLD III+IV, n = 66; mean ± SE, 6.23 ± 0.06) showing lower QKI levels than both non-COPD (n = 83; 6.42 ± 0.03) and mild-to-moderate COPD (GOLD I+II, n = 106; 6.43 ± 0.03) groups (P < 0.01 and P < 0.05, respectively) (Fig. 1A). QKI expression also correlated positively with lung function parameters, including percent predicted FEV_1_ (r = 0.15, P < 0.05; Fig. 1B) and FVC (r = 0.17, P < 0.01; Fig. 1C).

**Figure 1:**
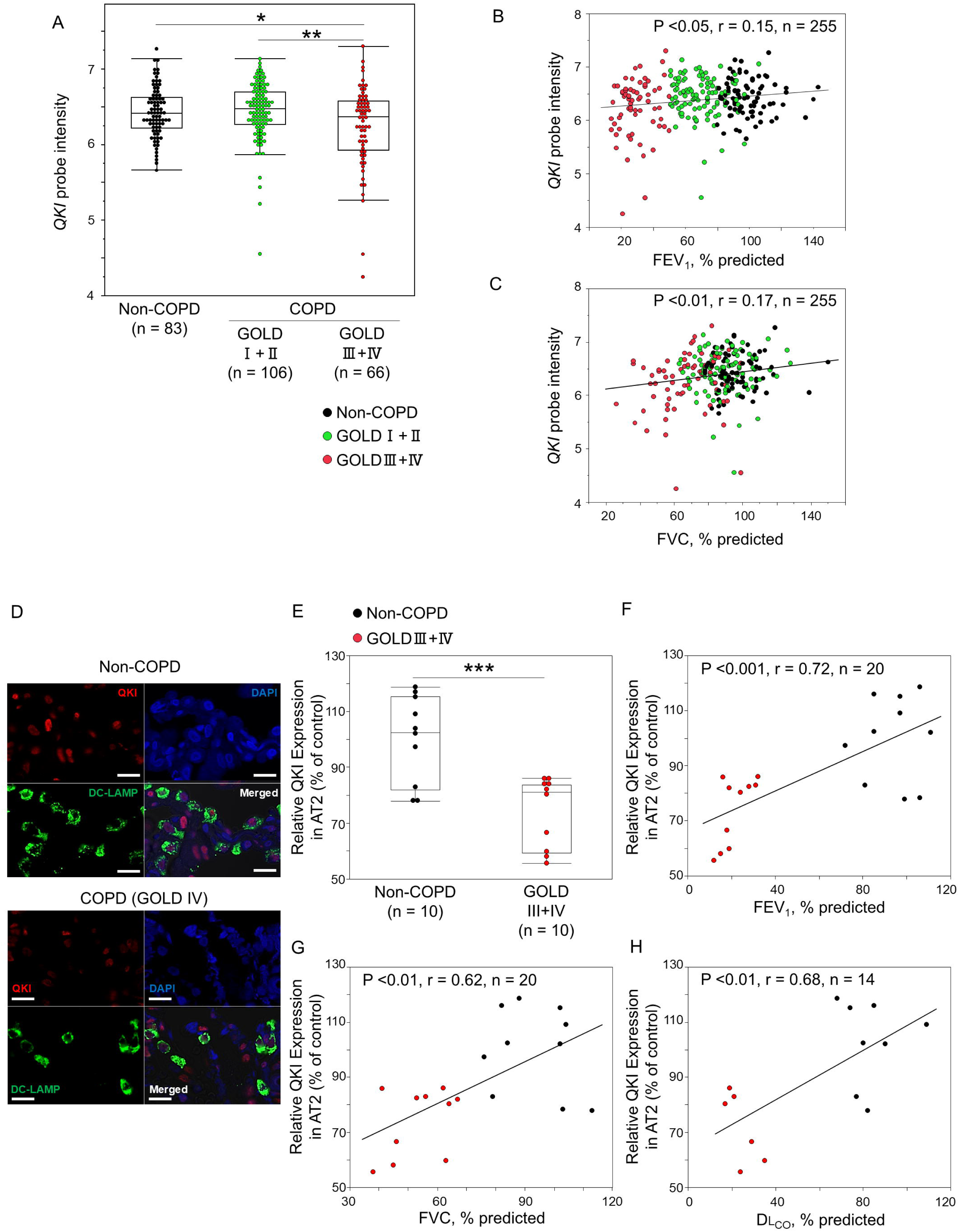
QKI Expression Is Reduced in COPD Lungs and Associated with Impaired Lung Function. (A-C) *QKI* gene expression levels were measured using a lung tissue microarray (*QKI* probe name: a_23_p81760). A total of 255 patients’ data from COPD patients and non-COPD patients with or without smoking history were analyzed. (A) *QKI* signal intensity was categorized into three groups: non-COPD (n = 83), mild to moderate COPD (GOLD stages I and II, n = 106), and severe to very severe COPD (GOLD stages III and IV, n = 66). Data are presented as box plots with individual data points and analyzed using one-way ANOVA followed by pairwise testing with the Tukey-Kramer HSD test. * Indicates P <0.05 compared to the non-COPD. ** Indicates P <0.01 compared to the GOLD I+II. (B, C) Correlation between QKI signal intensity and pulmonary function parameters: (B) the percentage predicted of post-bronchodilator FEV_1_ and (C) FVC. Statistical significance and correlation coefficients were determined using Pearson’s correlation. (D-H) QKI protein expression in AT2 cells was measured in lung sections by immunofluorescence staining. Non-COPD and severe to very severe COPD (GOLD III+IV) lungs were evaluated, and all samples were obtained from patients with a smoking history. At least fifty AT2 cells were evaluated from at least three different fields of view in each sample, and the average QKI fluorescence intensity was normalized to the non-COPD samples and calculated as relative QKI expression in AT2 cells (% of control) (also see online supplement). (D) Representative immunofluorescence images of lung sections from non-COPD and COPD (GOLD IV) stained for QKI (red), DC-LAMP (green; a marker of AT2 cells), and DAPI (blue). Scale bars, 50 um. (E) Comparison of relative QKI expressions in AT2 cells between non-COPD and severe and very severe COPD patients’ lungs. The data were analyzed using a student’s t test. *** Indicates P <0.001. (F-H) Correlation between relative QKI expressions in AT2 cells and pulmonary function parameters: (F) the percentage predicted of post-bronchodilator FEV_1_, (G) FVC, and (H) DL_CO_. Statistical significance and correlation coefficients were determined using Pearson’s correlation.

To assess alveolar epithelial cell-specific QKI expression, we performed immunofluorescence on FFPE sections from non-COPD (n = 10) and severe to very severe COPD (GOLD III+IV, n = 10) lungs (patient characteristics are described in Table E3). QKI was broadly expressed in several lung cell types, including alveolar epithelial cells, fibroblasts, and macrophages (Fig. 1D and Fig. E2). Notably, AT2 cells from patients with severe to very severe COPD had significantly lower QKI expression than those from non-COPD individuals (P < 0.001; Fig. 1E). AT2 cell-specific QKI expression correlated strongly with several lung function measures, including percent predicted FEV_1_ (r = 0.72, P < 0.001; Fig. 1F), FVC (r = 0.62, P < 0.01; Fig. 1G), and DL_CO_ (r = 0.68, P < 0.01; Fig. 1H). Collectively, these findings demonstrate reduced QKI expression in severe COPD and identify AT2 cells as an important site of QKI downregulation associated with impaired lung function.

### QKI Deficiency in Lung Epithelial Cells Distrupts Alveolar Homeostasis and Induces Emphysema in Mice

To elucidate QKI’s role in lung function, we generated LEC-specific QKI knockout mice (QKI^Δ/Δ^ mice) using *Sftpc*-Cre and *Qki^flox/flox^* mice. The Sftpc-Cre transgene used in this study is active in embryonic lung epithelial progenitors and mediates broad recombination throughout the pulmonary epithelium (26, 27). In *Qki^flox/flox^* control mice, QKI was broadly expressed in several lung cell types, including alveolar and bronchial epithelial cells, macrophages, endothelial cells, and mesenchymal cells. In contrast, QKI^Δ/Δ^ mice specifically lacked QKI in alveolar and bronchial epithelial cells (Fig. 2A and Fig. E3).

**Figure 2:**
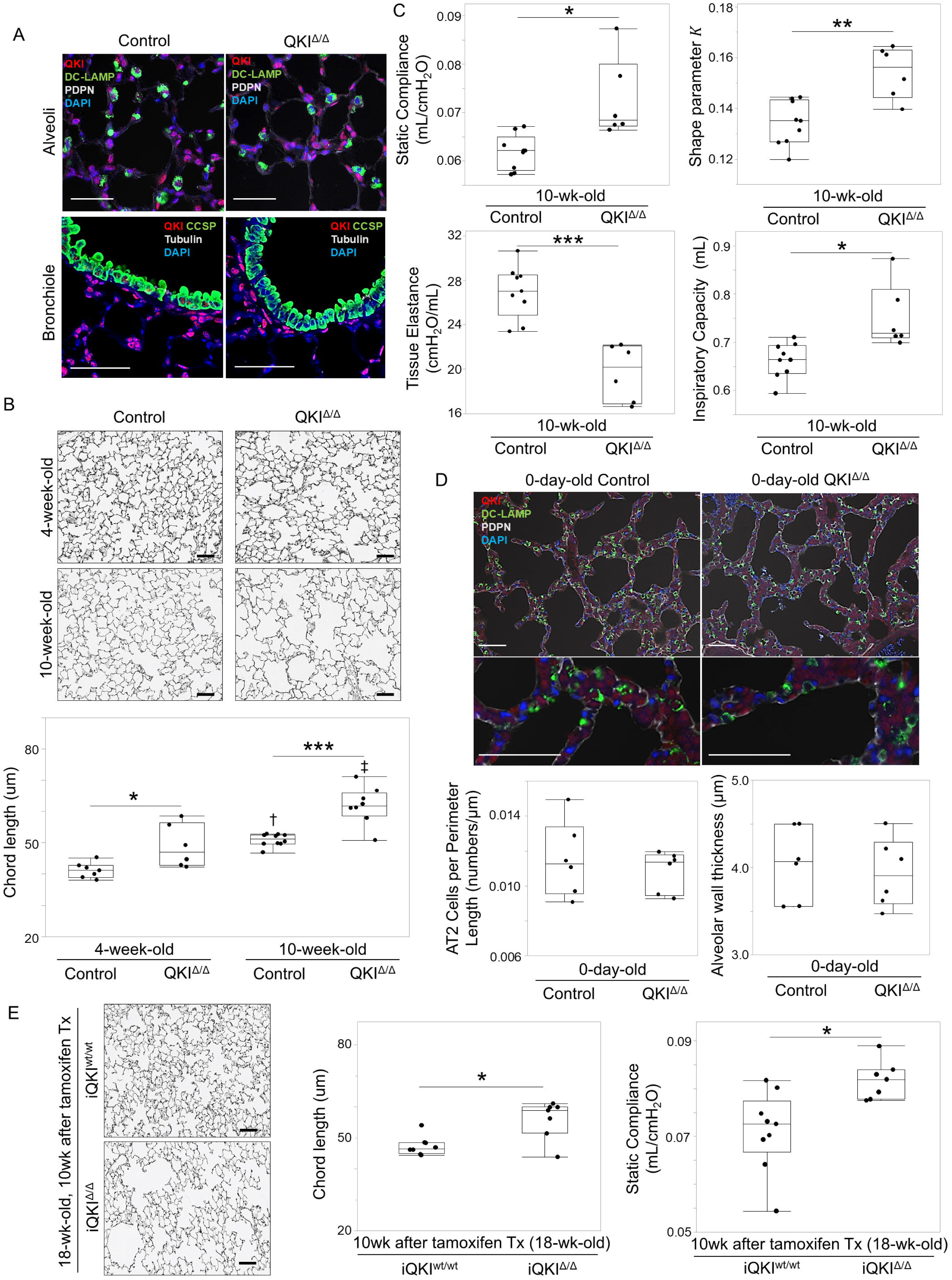
QKI Deficiency in Lung Epithelial Cells Disrupts Alveolar Homeostasis and Induces Emphysema in Mice. (A) Representative immunofluorescence images of lung sections from control and QKI^Δ/Δ^ mice stained for alveoli; QKI (red), DC-LAMP (green; a marker of alveolar type 2 epithelial cells), PDPN/Podoplanin (white; a marker of alveolar epithelial cells), and DAPI (blue) and bronchiole; QKI (red), CCSP/Club Cell Secretory Protein (green; a marker of club cells), acetylated tubulin (white; a marker of ciliated cells), and DAPI (blue). Scale bars, 50 um. (B) Representative hematoxylin and eosin (H&E) staining of lung sections from 4-week-old and 10-week-old control and QKI^Δ/Δ^ mice. Quantification of alveolar chord length was performed using a semi-automated method based on deep learning segmentation. n = 6-10 mice per group. (C) Pulmonary function parameters, including static compliance (Cst), shape parameter *K*, tissue elastance (H), and inspiratory capacity (IC) were measured using a FlexiVent system in 10-week-old control and QKI^Δ/Δ^ mice (n = 6-9 mice per group). (D) Immunofluorescence analysis of lung sections from newborn (0-day-old) control and QKI^Δ/Δ^ mice stained for QKI (red), DC-LAMP (green), and DAPI (blue). Quantification of the number of AT2 cells per alveolar perimeter length and alveolar wall thickness is shown. n = 6 mice per group. Scale bar, 50 μm. (E) Histological and physiological analysis of inducible QKI knockout mice (iQKI^Δ/Δ^), generated by administering tamoxifen to 8- to 9-week-old *Sftpc^tm1(cre/ERT2)Blh^*/ROSA26 *^flox/flox^* mice. Lung tissue was collected 10 weeks after induction (18-20 weeks old). Chord length and static compliance were measured. n = 7-9 mice per group. Box-and-whisker plots show individual data points. Statistical analyses were performed using one-way ANOVA followed by Tukey-Kramer HSD test for comparisons among multiple groups. Student’s t-test was used for two-group comparisons with equal variance; otherwise, Welch’s t-test was applied. * Indicates P <0.05. ** Indicates P <0.01. *** Indicates P <0.001. † Indicates P <0.01 compared to the same genotype at 4 weeks of age. ‡ Indicates P <0.0001 compared to the same genotype at 4 weeks of age.

First, we assessed alveolar structure by measuring alveolar chord length (CL) in four- and ten-week-old control and QKI^Δ/Δ^ mice (study design shown in Fig. E1). In QKI^Δ/Δ^ mice, CL was significantly greater than in controls at both ages, with a more pronounced difference at ten weeks (Fig. 2B). These findings indicate that loss of epithelial results in progressive airspace enlargement. Consistent with these structural alterations, static compliance (Cst) was elevated in QKI^Δ/Δ^ mice, indicating increased lung distensibility (Fig. 2C). This was supported by an increase in the shape parameter K, reflecting altered pressure–volume relationships. Tissue elastance (H), which inversely reflects tissue stiffness, was decreased, corroborating the loss of elastic recoil. Inspiratory capacity (IC) was also significantly increased, consistent with hyperinflation. Together, these changes capture the cardinal physiological features of emphysema. In addition, QKI^Δ/Δ^ mice exhibited increased numbers of lung inflammatory cells, primarily macrophages (Table E4), suggesting that QKI^Δ/Δ^ LECs may promote macrophage accumulation within the lung.

Next, to determine whether QKI loss has a greater impact on emphysema formation during embryonic development or during the postnatal period, we examined the lungs of QKI^Δ/Δ^ mice immediately after birth. In this model, *Sftpc*-Cre is activated at embryonic day 10.5 (E10.5) (27). The number of AT2 cells per alveolar perimeter length did not differ significantly between newborn (0-day-old) control and QKI^Δ/Δ^ mice. In addition, alveolar wall thickness showed no significant difference, suggesting that early alveolar structural organization remained intact despite QKI deficiency at this stage (Fig. 2D).

To examine the function of *QKI* in adult AT2, we generated a CreER-based model to delete QKI from AT2 cells after maturation (Fig. E1). When QKI was specifically deleted in mature AT2 cells (at 8–9 weeks of age), spontaneous emphysematous changes and elevated static lung compliance became apparent 10 weeks post-deletion (Fig. 2E). Taken together, these data indicate that QKI is dispensable for early alveolar patterning but is required for postnatal alveolar maintenance.

### QKI Deficiency Enhances Apoptosis and Reduces Colony Formation Efficiency in AT2 Cells

To characterize the phenotype of QKI^Δ/Δ^ AT2 cells, we conducted ex vivo three-dimensional spheroid assays to measure colony formation efficiency (CFE). QKI^Δ/Δ^ AT2 spheroids displayed significantly lower CFE than control AT2 spheroids (Fig. 3A). We then performed flow cytometric analyses using Ki-67 and Annexin V/PI staining. Although the proportion of Ki-67-positive AT2 cells remained unchanged, QKI^Δ/Δ^ AT2 spheroids had a higher proportion of Annexin V-positive cells (Fig. 3B). Notably, early apoptotic (Annexin V-positive/PI-negative) cells were significantly more abundant in QKI^Δ/Δ^ AT2 spheroids than in controls, suggesting that increased apoptosis, rather than decreased proliferation, accounts for the reduced CFE. Consistently, cleaved caspase-3 protein levels were elevated in isolated QKI^Δ/Δ^ AT2 cells compared with control cells (Fig. 3C). Together, these data indicate that QKI deficiency impairs AT2 cell CFE and is associated with increased apoptotic cell death.

**Figure 3:**
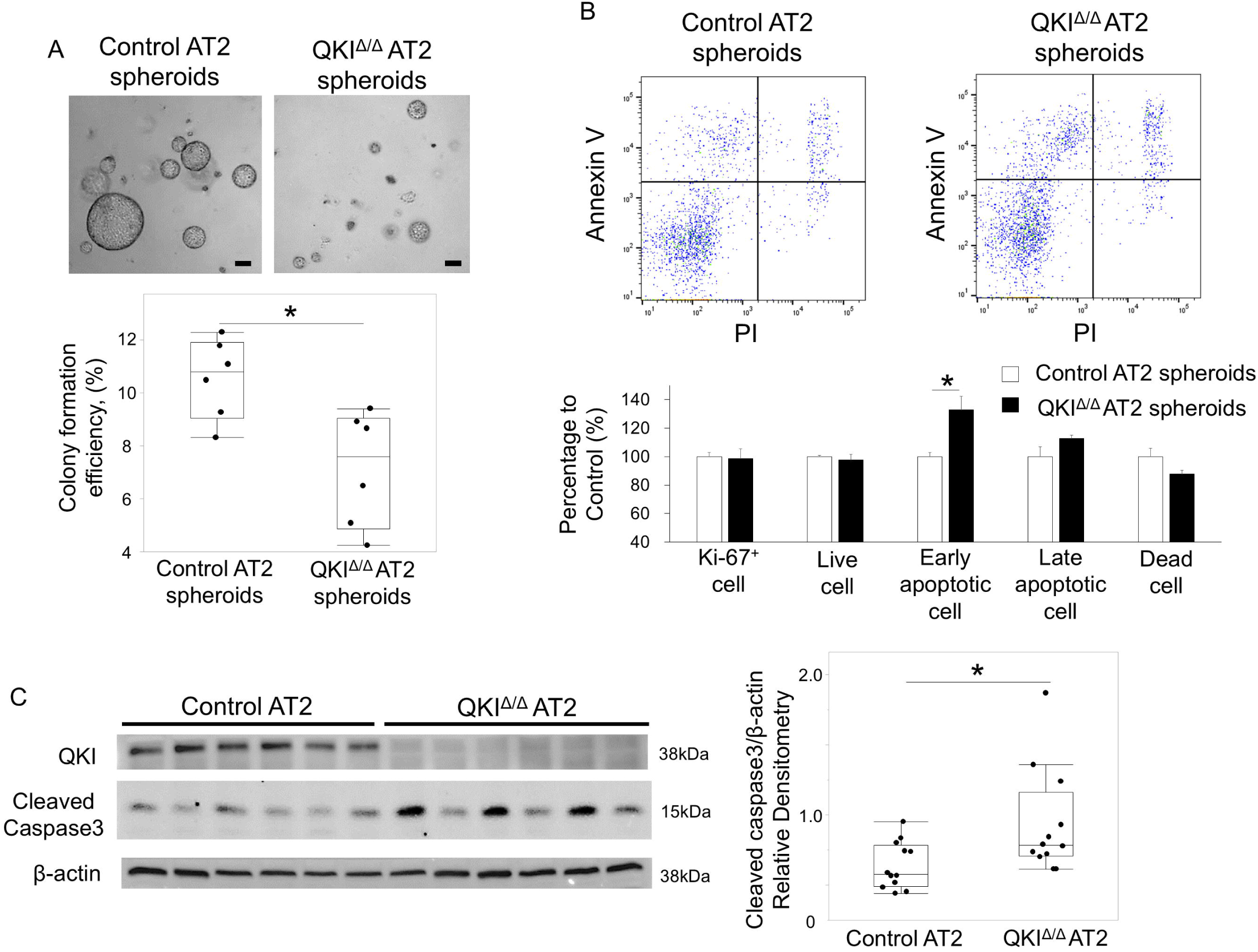
QKI Deficiency Enhances Apoptosis and Reduces Colony Formation Efficiency in AT2 Cells. (A) Spheroid assay of isolated AT2 cells from 10-week-old control and QKI^Δ/Δ^ mice. A total of 1,000 AT2 cells were embedded in Matrigel and cultured for 10 days. Representative images and quantification of colony formation efficiency are shown. Scale bar, 100 μm. n = 6 wells derived from two mice per group. (B) Flow cytometric analysis of AT2 cells from spheroids stained with Ki-67, Annexin V, and propidium iodide (PI). The proportions of Ki-67-positive cells and Annexin V-positive early apoptotic cells are shown. Cell fractions were determined by Annexin V and PI staining (also see online supplement). n = 6 mice per group. Bars indicate mean ± SEM. (C) Immunoblotting of isolated AT2 cells from control and QKI^Δ/Δ^ mice. QKI and cleaved caspase-3 protein levels were analyzed. Relative expression levels were quantified. n = 12 mice per group. Box-and-whisker plots show individual data points. Statistical analysis was performed using Student’s t-test. *P indicates < 0.05.

### QKI Deficiency Induces Transcriptome-Proteome Discordance in Mitochondrial and Metabolic Pathways

To define molecular programs altered by QKI loss in AT2 cells, we performed mRNA sequencing (RNA-seq) and proteomic profiling using independent control and QKI^Δ/Δ^ AT2 cell samples for each omics dataset (Fig. 4A). Principal component analysis and overview differential analyses confirmed group-level transcriptomic and proteomic changes in QKI^Δ/Δ^ AT2 cells (Fig. 4B and C; Fig. E5A and B).

**Figure 4:**
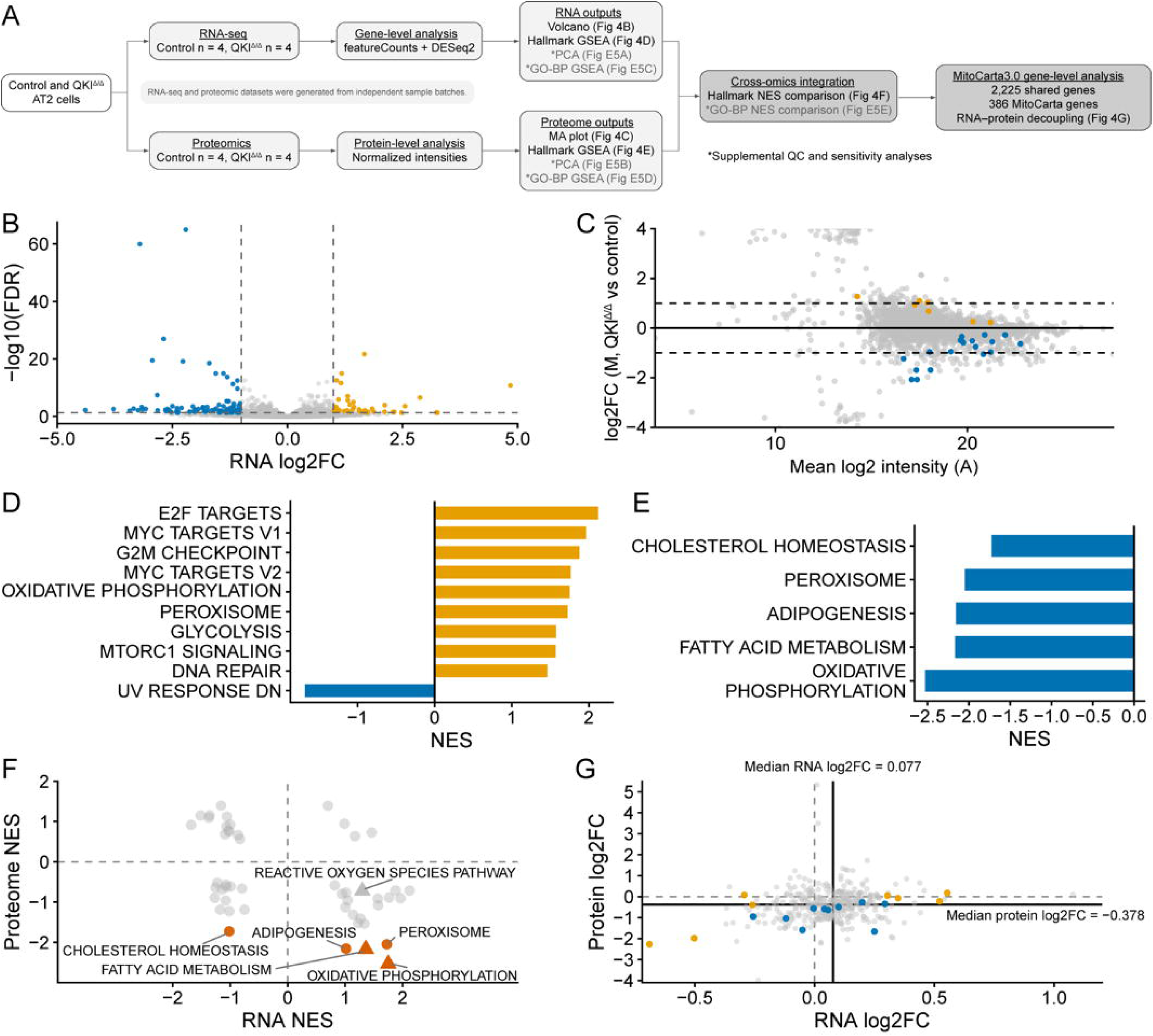
QKI Deficiency Induces Transcriptome-Proteome Discordance in Mitochondrial and Metabolic Pathways. (A) Schematic overview of the multi-omics analysis workflow. AT2 cells isolated from control and QKI^Δ/Δ^ mice were analyzed by RNA-seq and proteomics using independent sample sets for each omics dataset. (B) RNA-seq volcano plot showing differentially expressed genes in QKI^Δ/Δ^ versus control AT2 cells. All annotated genes with available log2FC and FDR values were included. Orange and blue points indicate genes significantly upregulated and downregulated in QKI^Δ/Δ^ cells, respectively, defined as FDR < 0.05 and |log2FC| ≥ 1. Dashed lines indicate the visualization thresholds. (C) Proteomic MA plot showing protein abundance changes in QKI^Δ/Δ^ versus control AT2 cells. A was calculated as the mean log2(normalized protein intensity + 1) across all samples, and M was calculated as the difference between the mean log2(normalized protein intensity + 1) in QKI^Δ/Δ^ and control samples. Orange and blue points indicate proteins upregulated and downregulated, respectively, in QKI^Δ/Δ^ cells with FDR < 0.05. For visualization, the y-axis is displayed from −4 to 4. (D) Hallmark GSEA of RNA-seq data using a protein-coding ranked gene list. Bars indicate normalized enrichment scores (NES) for Hallmark pathways significantly enriched in QKI^Δ/Δ^ versus control AT2 cells. (E) Hallmark GSEA of proteomic data. Bars indicate NES for significantly enriched Hallmark pathways in QKI^Δ/Δ^ versus control AT2 cells. Mitochondrial and metabolic pathways, including oxidative phosphorylation and fatty acid metabolism, showed negative enrichment at the protein level. (F) Integrated comparison of RNA-seq and proteomic Hallmark GSEA results. Each point represents one Hallmark gene set. Point color denotes proteomic GSEA significance: vermilion, proteome FDR < 0.05; gray, proteome FDR ≥ 0.05. Point shape denotes pathway class: triangle, mitochondria-related Hallmark pathway; circle, other Hallmark pathway. Mitochondria-related Hallmark pathways were defined as oxidative phosphorylation, fatty acid metabolism, and reactive oxygen species pathway. (G) Gene-level RNA–protein comparison of MitoCarta3.0 genes detected in both RNA-seq and proteomic datasets. Each point represents one shared MitoCarta3.0 gene. Points are colored by significance class: vermilion, significant in both datasets (no genes fell into this category in this dataset); orange, significant only in RNA-seq; blue, significant only in proteomics; and gray, not significant in either dataset. Dashed lines indicate zero log2FC, and solid lines indicate median RNA log2FC and median protein log2FC.

Hallmark Gene Set Enrichment Analysis (GSEA) revealed positive RNA-level enrichment of oxidative phosphorylation, peroxisome, and glycolysis (Fig. 4D). In contrast, proteomic Hallmark GSEA showed negative enrichment of mitochondrial and metabolic pathways, including oxidative phosphorylation, fatty acid metabolism, adipogenesis, peroxisome, and cholesterol homeostasis (Fig. 4E). RNA-seq GSEA also revealed enrichment of MYC targets, E2F targets, and G2M checkpoint gene sets, consistent with activation of proliferative or stress-associated transcriptional programs in QKI^Δ/Δ^ AT2 cells. These pathways, however, were not enriched at the proteome-level pathway output. Because this analysis was designed to detect coordinated pathway-level changes, individual gene-level results were not emphasized in the main figure but are provided in the Supporting Data Values and Supplementary Tables, including core enriched genes for GSEA terms. Direct comparison of Hallmark Normalized Enrichment Score (NES) values highlighted RNA–protein discordance among mitochondrial and metabolic pathways (Fig. 4F).

We next examined this discordance at the gene level using MitoCarta3.0. Among 386 MitoCarta3.0 genes detected in both datasets, median RNA log_2_FC was 0.077, whereas median proteome log_2_FC was -0.378. The RNA-protein discrepancy was significantly shifted above zero (median RNA log_2_FC − protein log_2_FC = 0.46; Wilcoxon signed-rank test, P < 2.2×10^−16^), consistent with mitochondrial RNA-protein decoupling in QKI^Δ/Δ^ AT2 cells (Fig. 4G). Gene Ontology Biological Process (GO-BP) GSEA and GO-BP NES integration further supported this mitochondrial/metabolic transcriptome–proteome discordance (Fig. E5C-E). Together, these findings demonstrate significant transcriptome-proteome discordance in QKI^Δ/Δ^ AT2 cells, particularly in mitochondrial and metabolic pathways, suggesting that QKI is required for coordinated RNA and protein expression.

### QKI Deficiency Produces Respiratory Protein-Deficient Mitochondria

To investigate the association between QKI and mitochondrial function, we generated QKI^−/−^ BEAS-2B (B2B) cells using CRISPR/Cas9 (Fig. E4). Although BEAS-2B cells are not alveolar epithelial cells, they provide a tractable human epithelial system to test cell-intrinsic consequences of QKI loss and reversibility. Real-time mitochondrial oxidative respiration in QKI^−/−^ B2B cells was measured using the Seahorse Mito Stress Test. QKI^−/−^ cells had a significantly reduced oxygen consumption rate (OCR) compared with control cells, with reduced maximal respiration (P < 0.01; mean ± SE, control: 108.5 ± 11.2 vs. QKI^−/−^ B2B: 48.1 ± 11.8 pmol/min/normalized unit), indicating compromised mitochondrial respiratory capacity (Fig. 5A). In addition, QKI^−/−^ cells showed a significant increase in basal extracellular acidification rate (P < 0.01; mean ± SE, control: 22.2 ± 2.5 vs. QKI^−/−^ B2B: 37.3 ± 2.6 pmol/min/normalized unit), consistent with enhanced reliance on glycolytic metabolism. Cellular energy metabolism analysis revealed no significant difference in mitochondrial ATP production in QKI^−/−^ cells, indicating a shift in cellular energy metabolism toward glycolysis (Fig. 5B).

**Figure 5:**
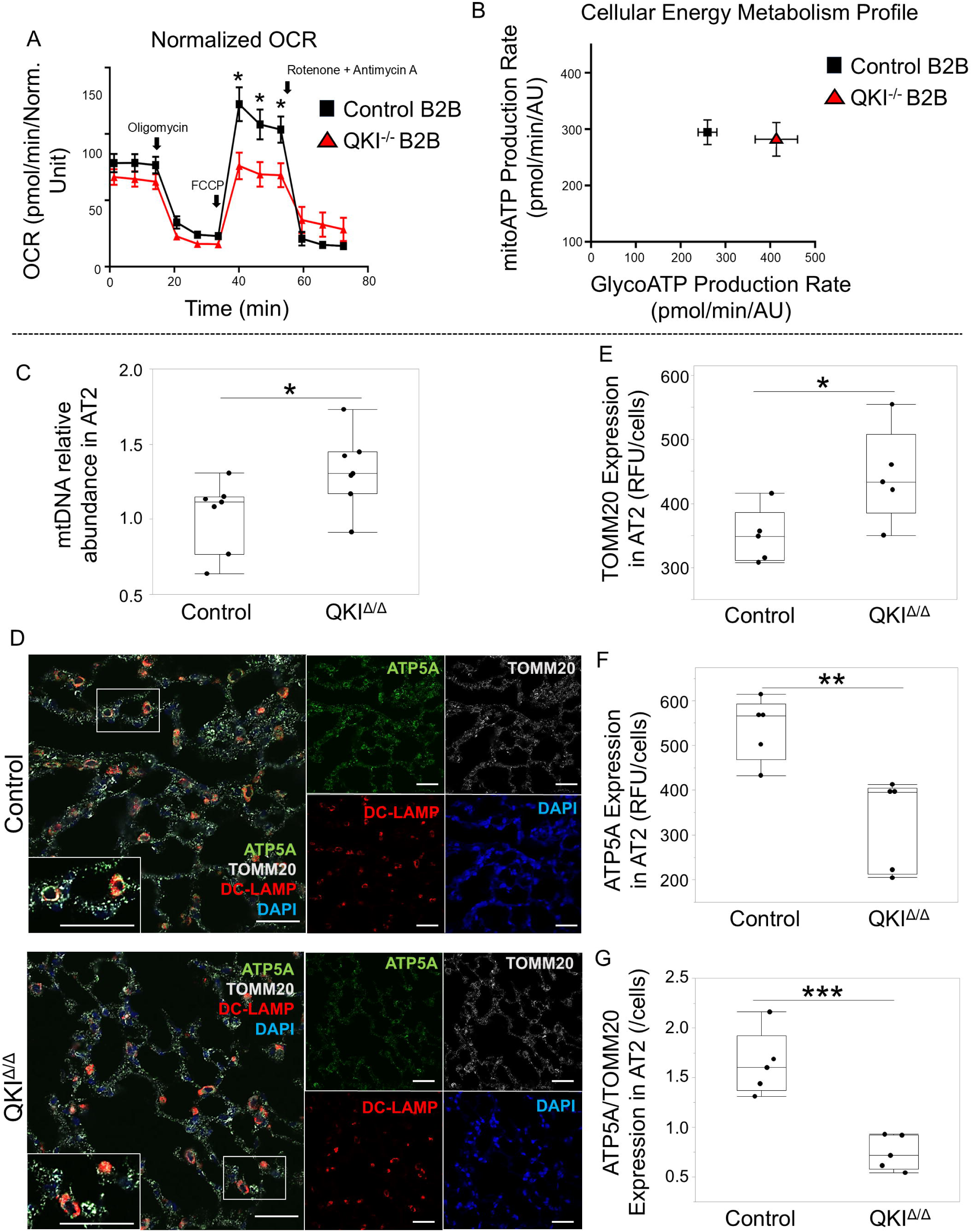
QKI Deficiency Produces Respiratory Protein-Deficient Mitochondria. (A, B) Seahorse-based metabolic analyses were performed in control and QKI^⁻/⁻^ Beas 2B (B2B) cells. (C–G) Mitochondrial abundance and protein expression were assessed in isolated murine AT2 cells and lung sections from control and QKI^Δ/Δ^ mice. (A) Oxygen consumption rate (OCR) was measured using the Seahorse XF Analyzer in control and QKI^⁻/⁻^ B2B cells following sequential treatment with oligomycin, FCCP, and rotenone/antimycin A. OCR was normalized to baseline. n = 10 control and 9 QKI^⁻/⁻^ B2B cell wells. (B) Cellular energy metabolism profile showing mitoATP production (oxidative phosphorylation) and glycoATP production (glycolysis). (C) Relative abundance of mitochondrial DNA (mtDNA) in isolated AT2 cells determined by qPCR (n = 7 mice per group). (D) Representative immunofluorescence images of lung sections from Control and QKI^Δ/Δ^ mice stained for ATP5A (green), TOMM20 (white), DC-LAMP (red; AT2 cell marker), and DAPI (blue; nuclei). Scale bars: 50 um. (E–G) Quantification of fluorescence intensity for TOMM20 (E), ATP5A (F), and the ratio of ATP5A/TOMM20 expression (G) in AT2 cells (n = 5 mice per group). Box-and-whisker plots show individual data points. Student’s t-test was used when variance and sample size were equal; otherwise, Welch’s t-test was applied. *P < 0.05, **P < 0.01, ***P < 0.001.

To determine whether this metabolic remodeling is accompanied by altered mitochondrial abundance and functional protein composition in a disease-relevant epithelial context, we next assessed mitochondrial content and respiratory-chain markers in murine QKI^Δ/Δ^ AT2 cells. In QKI^Δ/Δ^ AT2 cells, qPCR analysis demonstrated a significant increase in mitochondrial DNA (mtDNA) abundance (Fig. 5C), and immunofluorescence confirmed an increase in the outer membrane marker TOMM20 (Fig. 5D and E). However, despite these compensatory signals, expression of the functional inner membrane protein ATP5A was markedly reduced (Fig. 5D and F), resulting in a significant decline in the ATP5A/TOMM20 ratio (Fig. 5G) in QKI^Δ/Δ^ AT2 cells.

Together, these findings indicate that QKI deficiency is associated with mitochondrial dysfunction and the accumulation of respiratory protein-deficient mitochondria.

### Re-expression of QKI Partially Rescues Cellular and Mitochondrial Dysfunction in QKI-deficient Cells

We next investigated whether QKI loss-associated phenotypes could be rescued by restoring QKI expression. For this experiment, we generated QKI add-back B2B cells by transducing QKI^−/−^ B2B cells with a QKI expression plasmid (Fig. E4).

Consistent with findings in QKI^Δ/Δ^ murine AT2 cells, QKI^−/−^ B2B cells exhibited reduced CFE capacity (Fig. 6A), reduced MTT-based metabolic activity (Fig. 6B), and increased apoptosis (Fig. 6C). QKI^−/−^ B2B cells also exhibited increased mitochondrial ROS levels (Fig. 6D) and elevated mitochondrial membrane potential (Fig. 6E), consistent with mitochondrial dysfunction. Re-expression of QKI in QKI add-back B2B cells partially reversed these abnormalities, restoring CFE (Fig. 6A) and metabolic activity (Fig. 6B) and attenuating apoptosis (Fig. 6C), mitochondrial ROS (Fig. 6D), and mitochondrial membrane potential (Fig. 6E) toward control levels.

**Figure 6:**
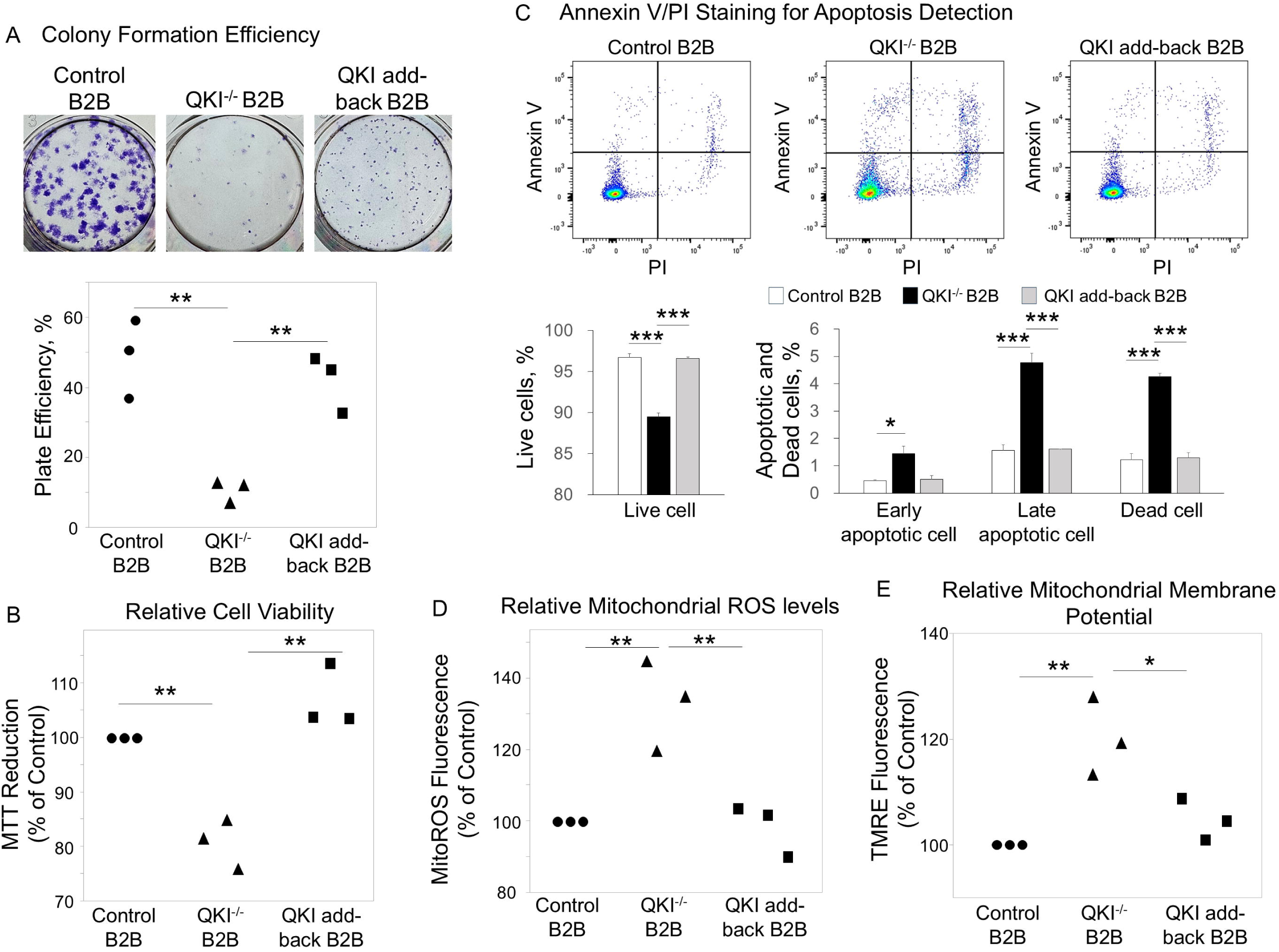
Re-expression of QKI Partially Rescues Cellular and Mitochondrial Dysfunction in QKI-deficient Cells. (A) Clonogenic assay of Beas 2B (B2B) cells. Two hundred cells were cultured for 10 days, and colony formation efficiency (plate efficiency) was quantified. Each data point represents the average of two technical replicates from one biological replicate. A total of three independent experiments were performed (n = 3 biological replicates). Representative images and quantification are shown. (B) Metabolic activity was assessed by MTT (3-(4,5-dimethylthiazol-2-yl)-2,5-diphenyltetrazolium bromide) assay. Each data point represents the average of three wells per experiment, with three independent experiments performed (n = 3 biological replicates). MTT reduction is shown as a percentage relative to control B2B cells. (C) Apoptosis was analyzed by flow cytometry using Annexin V and Propidium Iodine staining. Each data point represents a single well, with three independent experiments performed (n = 3 biological replicates). The bar chart indicates the mean and the standard error of the mean (SEM). (D) Mitochondrial reactive oxygen species (ROS) levels were measured by MitoROS fluorescence and normalized to control B2B cells. Each data point represents the average of six wells per experiment, with three independent experiments performed (n = 3 biological replicates). (E) Mitochondrial membrane potential was assessed using tetramethylrhodamine ethyl ester (TMRE) fluorescence normalized to control B2B cells. Each data point represents the average of six wells per experiment, with three independent experiments performed (n = 3 biological replicates). Statistical analysis was performed using one-way ANOVA with Tukey-Kramer HSD test for multiple group comparisons. *P < 0.05, **P < 0.01, ***P < 0.001, ****P < 0.0001.

**Figure 7:**
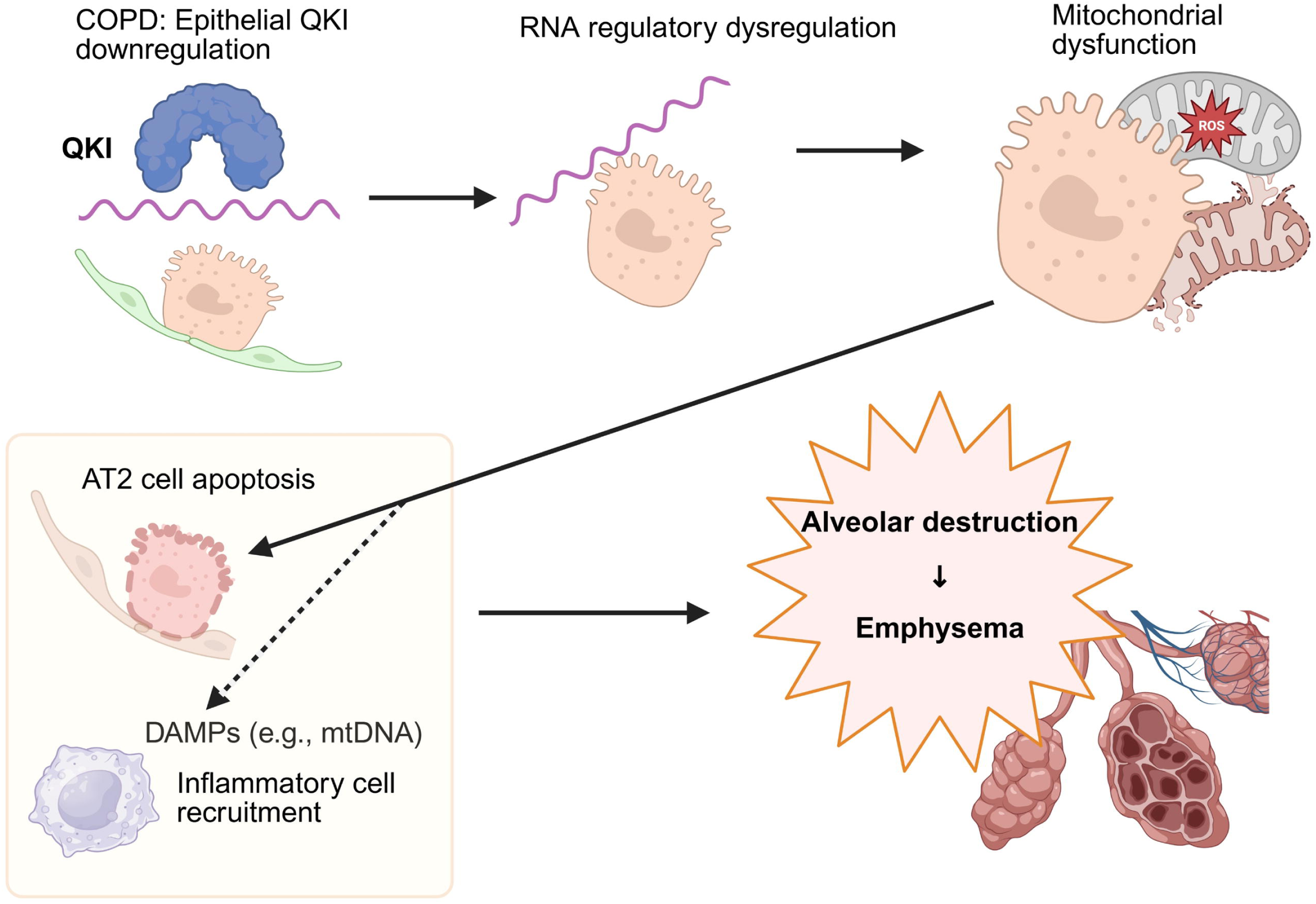
Proposed Model linking Epithelial QKI Downregulation to Mitochondrial dysfunction and Emphysematous Remodeling in COPD. Reduced QKI expression in alveolar type 2 epithelial (AT2) cells in COPD is hypothesized to cause dysregulation of RNA regulatory programs, leading to mitochondrial dysfunction. Mitochondrial injury promotes AT2 cell apoptosis and the release of damage-associated molecular patterns (DAMPs), including mitochondrial DNA (mtDNA), which may further drive inflammatory cell recruitment. These epithelial-intrinsic and inflammation-associated processes are proposed to converge on progressive alveolar destruction and emphysema. *This figure was Created with* BioRender.com.

These data suggest that QKI loss-induced dysfunction is at least partially reversible, supporting a crucial role for QKI in maintaining epithelial homeostasis.

## Discussion

Our study demonstrates that QKI, an RNA-binding protein previously implicated in cell differentiation and homeostasis (15, 21, 28), plays a critical role in maintaining alveolar integrity and lung function. In human COPD lungs, QKI expression, particularly in AT2 cells, was reduced and correlated with disease severity, including lower percent predicted FEV_1_ and DL_CO_ values. Consistently, mice with QKI deletion in LECs (QKI^Δ/Δ^) developed emphysematous remodeling characterized by airspace enlargement and increased static compliance, underscoring QKI’s essential role in maintaining alveolar homeostasis.

A central mechanistic implication of our work is that QKI loss compromises mitochondrial homeostasis, resulting in the accumulation of structurally present but functionally impaired mitochondria. QKI^Δ/Δ^ AT2 cells exhibited reduced colony formation efficiency and increased apoptosis in three-dimensional spheroid assays, and proteomic profiling revealed broad suppression of mitochondrial antioxidant and metabolic proteins. Importantly, the additional mitochondrial assays introduced in this revision strengthen the conclusion that QKI deficiency leads to the accumulation of dysfunctional mitochondria: mtDNA abundance and the outer membrane marker TOMM20 were increased, whereas the inner membrane oxidative phosphorylation component ATP5A was decreased, resulting in a markedly reduced ATP5A/TOMM20 ratio. These findings indicate that compensatory mitochondrial biogenesis is induced but fails to restore functional respiratory chain capacity. Consistently, human epithelial QKI^−/−^ cells showed impaired oxidative respiration, increased mitochondrial ROS, and hyperpolarized mitochondrial membrane potential, and these abnormalities were partially reversed by QKI add-back, supporting a cell-intrinsic and at least partially reversible mitochondrial phenotype. Increased apoptosis was a consistent feature of QKI loss across model systems, observed both in vivo in QKI^Δ/Δ^ AT2 cells (elevated Annexin V positivity and cleaved caspase-3) and in vitro in human QKI^−/−^ B2B cells, and it was attenuated by QKI re-expression. This concordance between the primary murine AT2 phenotype and the tractable human epithelial system indicates that apoptosis is a reproducible, cell-intrinsic consequence of QKI deficiency rather than a model-specific artifact, consistent with the increased epithelial apoptosis reported in COPD lungs. Notably, whereas QKI add-back restored the impaired phenotype in QKI-deficient BEAS-2B cells, pharmacologic interventions targeting oxidative stress and mitochondrial metabolism, including N-acetylcysteine, MitoQ (mitoquinone mesylate), nicotinamide mononucleotide (NMN), and 5-aminoimidazole-4-carboxamide-1-β-D-ribofuranoside (AICAR), did not produce measurable functional recovery (data not shown). These negative results argue against the possibility that the phenotype is driven solely by excess ROS or a single, readily correctable metabolic insufficiency. Instead, these findings support a model in which QKI contributes to broader RNA regulatory programs required for mitochondrial homeostasis and respiratory-chain integrity, such that supplementing redox capacity or activating generic metabolic pathways is insufficient unless the upstream QKI-dependent regulatory network is restored. Notably, these agents were evaluated as single-agent interventions in the absence of QKI; we therefore cannot exclude the possibility that they act as supportive cofactors that enhance the QKI add-back response or that combinatorial approaches are required. Whether such interventions augment, rather than replace, QKI-dependent restoration of mitochondrial function, and which functional protein components are most rate-limiting for rescue, will require dedicated combinatorial and gene-restoration experiments in future studies.

This interpretation is reinforced by the discordance between transcriptomic and proteomic profiles. While multiple mitochondrial proteins were reduced, mRNA sequencing identified enrichment of mitochondria-related Hallmark gene sets, including oxidative phosphorylation, peroxisome, and fatty acid metabolism, consistent with an attempted compensatory transcriptional response. Thus, the primary defect appears to lie downstream of transcriptional activation, and compensatory mitochondrial biogenesis signals are insufficient to overcome loss of functional protein components.

Beyond mitochondrial and metabolic pathways, RNA-seq also showed enrichment of growth/proliferative and activation-like gene sets, including MYC targets, E2F targets, G2M checkpoint, and MTORC1 signaling. Together with preserved Ki-67 staining, these findings suggest that QKI-deficient AT2 cells retain compensatory or activation-like responses at the transcriptomic and cellular levels. However, these pathways were not enriched at the proteome level. In the context of mitochondrial/metabolic protein depletion, this pattern suggests that activation-like responses may lack sufficient proteome-level metabolic support.

Mechanistically, QKI is well established to regulate RNA stability, alternative splicing, localization, and translation. Prior work indicates that QKI can influence metabolic homeostasis and interact with mitochondrial RNA (29), suggesting the possibility that QKI coordinates post-transcriptional expression of nuclear-encoded mitochondrial transcripts and/or their translation on mitochondrial-associated ribosomes. In this framework, QKI loss would uncouple transcriptional compensation from protein output, explaining the observed pattern of increased mitochondrial content markers alongside loss of key enzymatic and respiratory chain proteins. Future studies will be needed to define the precise co- and/or post-transcriptional mechanisms linking QKI deficiency to impaired mitochondrial function. Furthermore, recent human studies demonstrate that QKI regulates post-transcriptional expression of protective genes such as *SERPINA1* via alternative polyadenylation, supporting the concept that QKI-dependent RNA regulation is disrupted in COPD (30).

To distinguish developmental from maintenance functions, we analyzed lungs at birth (day 0), day 28, and 10 weeks of age. Although Sftpc-Cre–mediated QKI deletion begins at approximately E10.5 (27), QKI^Δ/Δ^ mice did not exhibit neonatal respiratory distress or overt structural abnormalities at birth. However, airspace enlargement was evident by four weeks and persisted through 10 weeks, indicating that QKI is dispensable for late-stage morphogenesis after Sftpc-Cre activation but becomes essential for postnatal alveolar maintenance. Moreover, inducible deletion in mature AT2 cells produced emphysematous changes within 10 weeks after deletion, underscoring an ongoing requirement for QKI in adult alveolar integrity. Future studies employing earlier-acting lung lineage drivers (e.g., Shh, Nkx2.1) (31, 32) could further delineate potential roles for QKI at earlier embryonic stages. A plausible explanation for this temporal selectivity is that early alveolar patterning is driven by redundant morphogenetic programs and a low biosynthetic and respiratory demand, whereas postnatal alveologenesis and subsequent maintenance impose a sustained requirement for high mitochondrial output to support surfactant synthesis, AT2 self-renewal, and repair. In this setting, QKI-dependent RNA regulatory programs that support mitochondrial function becomes rate-limiting, so that its loss is tolerated during development but progressively unmasks a bioenergetic deficit once AT2 cells must continuously sustain alveolar homeostasis under postnatal metabolic and oxidative load.

Because COPD pathogenesis involves complex epithelial–immune interactions and macrophages exhibit substantial QKI expression (33), we also evaluated myeloid-specific deletion (Lyz2-QKI^Δ/Δ^ mice, Fig. E6). In contrast to epithelial deletion, Lyz2-QKI^Δ/Δ^ mice did not show airspace enlargement or altered lung mechanics at 10 weeks of age. Although longer-term observation and stress models (e.g., cigarette smoke exposure, infectious insults, or aging) will be required to fully define the contribution of myeloid QKI, these baseline data suggest that AT2 cells, a key for surfactant production and epithelial repair, are particularly susceptible to QKI loss under homeostatic conditions.

From a pathophysiological perspective, our findings are consistent with extensive evidence that mitochondrial dysfunction is central to COPD (34, 35), promoting oxidative stress (36, 37), amplifying inflammatory signaling (38, 39), and activating cell death pathways (40). In QKI-deficient AT2 cells, decreased levels of mitochondrial antioxidant enzymes (Table 1, e.g., SOD2) and metabolic enzymes (e.g., PDHX and DLAT) provide plausible molecular substrates for increased ROS generation, impaired ATP production, and apoptosis. These mechanisms align with established models in which oxidative stress and epithelial injury drive progressive alveolar destruction (41–43). Our findings provide mechanistic insights into why certain patients may be more vulnerable to emphysematous changes due to impaired mitochondrial integrity and epithelial viability.

**Table 1.**
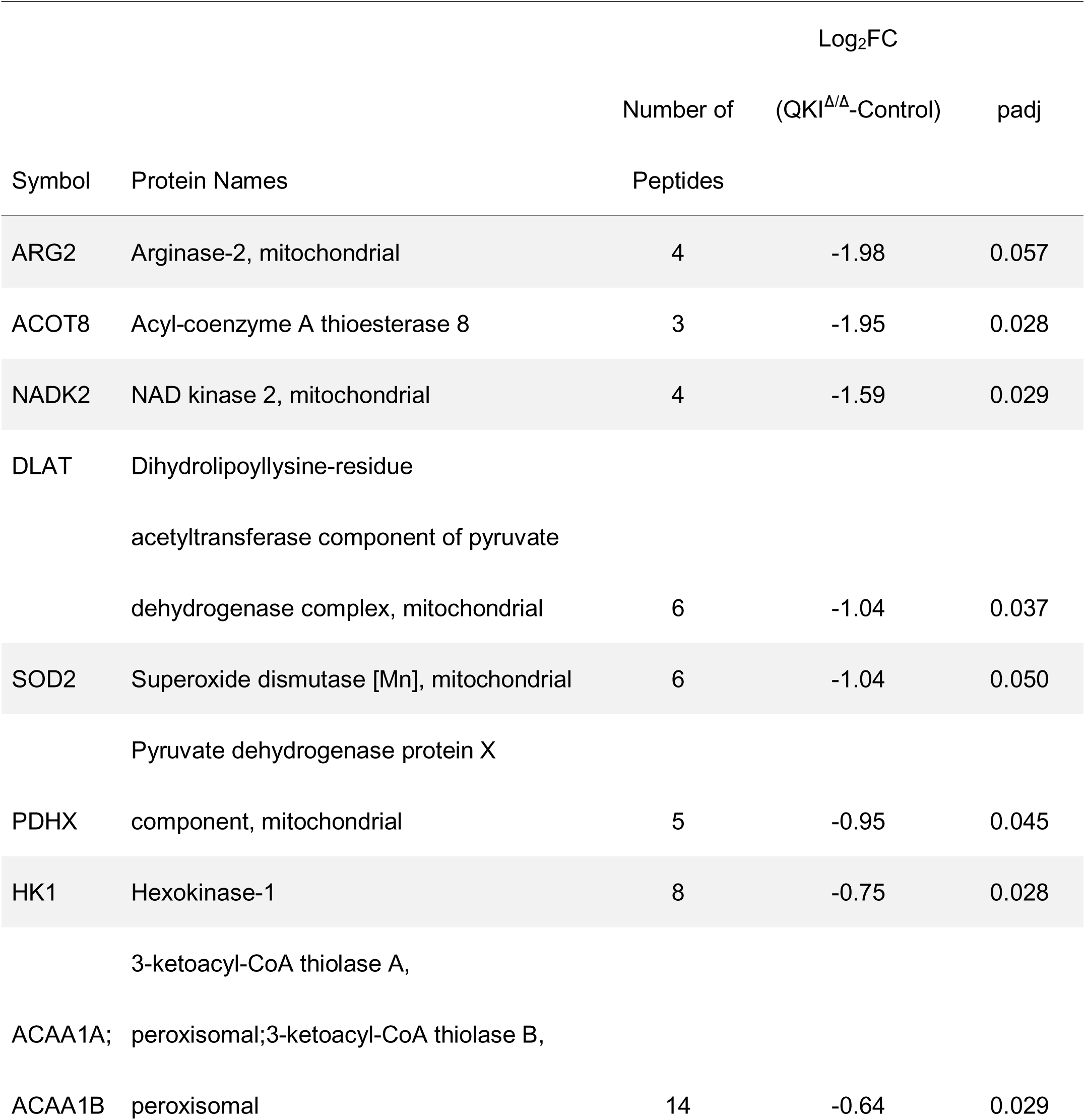
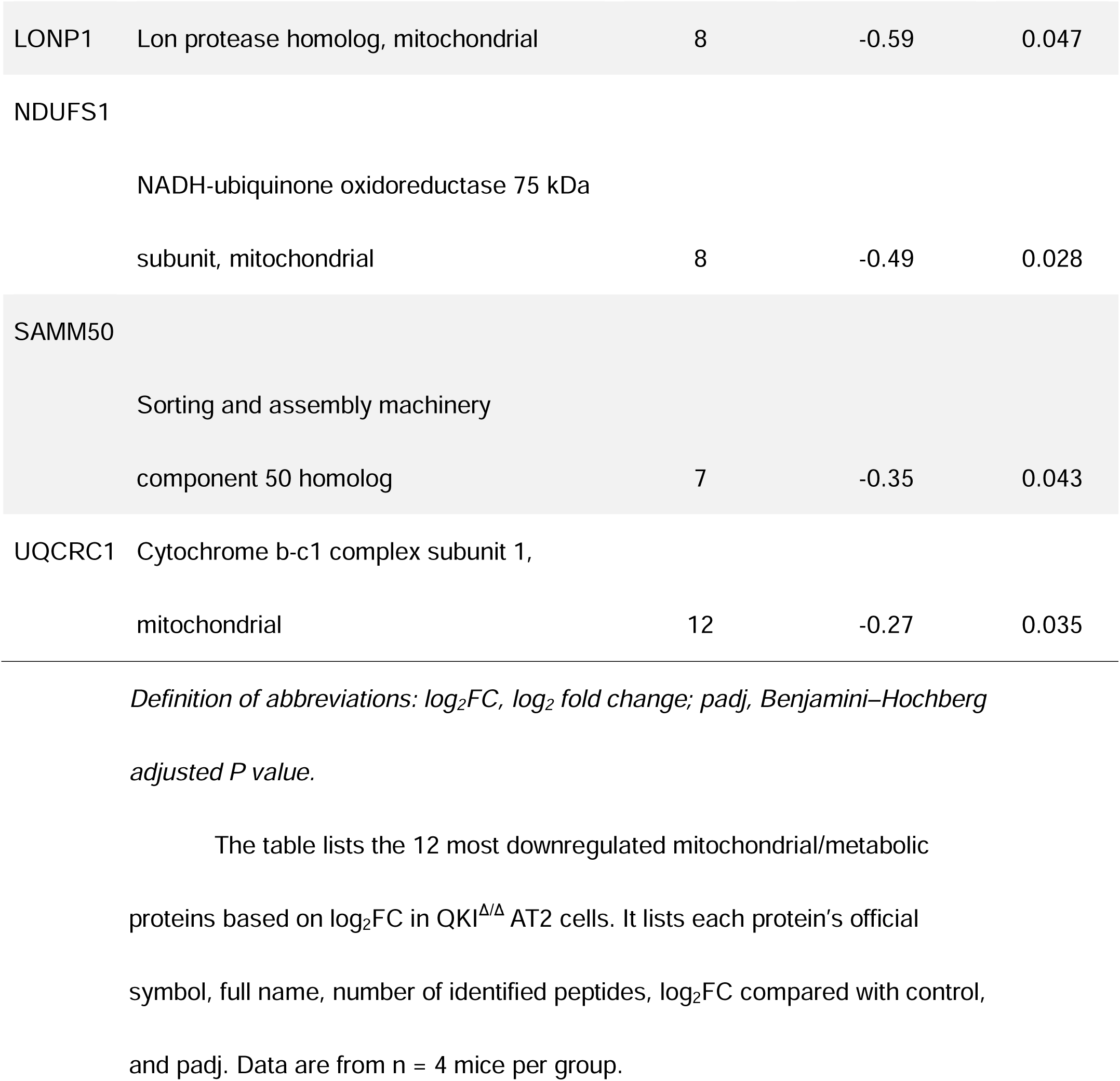
The 12 most downregulated mitochondrial proteins (based on Log_2_FC) in QKI^Δ/Δ^ mice AT2 cells.

Our study also expands the conceptual landscape of RNA-binding proteins in COPD. Prior work has largely focused on RBPs that regulate oxidative stress–related genes, senescence-associated secretory phenotypes, and inflammatory responses via mRNA turnover (44–48). By contrast, our data implicates QKI as a key regulator of mitochondrial homeostasis and epithelial viability, thereby providing a direct mechanistic link between RBP dysregulation, mitochondrial failure, and emphysematous remodeling.

Several limitations merit consideration. First, QKI is expressed across epithelial, endothelial, and immune compartments (12), and the relative contribution of each population may vary with disease stage and environmental exposures. Second, while our multi-omics analyses and mitochondrial markers support a model of dysfunctional mitochondrial accumulation, downstream quality-control processes, such as mitophagy, fission–fusion dynamics, and mitochondria–lysosome crosstalk, were not directly interrogated and should be addressed in future work. Third, our models do not fully recapitulate the systemic and exposure-related features of COPD, including accelerated aging and comorbidities; therefore, testing QKI-deficient mice in cigarette smoke or other injury models will be important to assess generalizability and to define context-dependent effects. Fourth, because QKI is an established regulator of lipid homeostasis and AT2 cells synthesize all subclasses of surfactant lipids, our study did not resolve whether QKI loss alters surfactant lipid synthesis in AT2 cells or perturbs the lipofibroblast–AT2 cell lipid-transfer axis that supports surfactant production; defining how QKI-dependent lipid metabolism intersects with the mitochondrial defect described here, in both AT2 cells and neighboring lipofibroblasts, will be an important direction for future work.

Despite these considerations, the strong association between QKI levels in human AT2 cells and pulmonary function, together with definitive mechanistic evidence from epithelial knockout models, highlights QKI as a potential therapeutic entry point. Current COPD therapies primarily relieve airflow limitation without preventing progressive alveolar destruction. Targeting post-transcriptional programs controlled by QKI, or downstream pathways that preserve mitochondrial quality and epithelial survival, could represent a complementary strategy aimed at maintaining alveolar structure. Although tissue-specific delivery remains a challenge, advances in RNA-based therapeutics and targeted pulmonary delivery platforms may enable approaches that augment QKI function or restore critical QKI-regulated pathways. We do not propose that mitochondrial dysfunction arises from a single downstream target of QKI, but rather from coordinated disruption of RNA regulatory programs governing mitochondrial homeostasis.

In conclusion, our study identifies QKI as an indispensable regulator of alveolar epithelial homeostasis. Loss of QKI in lung epithelial cells leads to emphysematous remodeling associated with mitochondrial dysfunction and apoptosis, likely driven by RNA regulatory mechanisms that uncouple compensatory transcriptional responses from mitochondrial protein integrity. These findings provide a mechanistic framework linking reduced QKI expression in human COPD to impaired alveolar maintenance and suggest QKI-centered pathways as candidates for disease-modifying intervention. Together, these findings support a model in which epithelial QKI preserves mitochondrial integrity and stress resilience, and its loss lowers the threshold for epithelial failure, leading to emphysema.

## Methods

### Human studies

#### Sex as a biological variable

Both male and female participants were included in the human analyses; the numbers of males and females are reported in Table E2 and E3.

### Analysis of QKI characteristics in human lung tissues and clinical data

Lung tissue microarray data and pulmonary function measurements were obtained from the Lung Tissue Research Consortium (LTRC). Formalin-fixed, paraffin-embedded (FFPE) lung tissue sections with corresponding clinical data were acquired from the NIH.

Lung tissue sections were immunostained as previously described (49), and images were acquired using a Nikon Ti microscope (Nikon Instruments Inc., Melville, NY, USA). Mean QKI fluorescence intensities in AT2 epithelial cells, identified by DC-LAMP (50), was quantified using ImageJ (National Institutes of Health, Bethesda, MD, USA). Expression levels were normalized by dividing each sample’s QKI expression by the average QKI expression of control samples. Detailed protocols for tissue processing and image analysis are provided in the online supplement.

### *In vitro* study with human bronchial epithelial cells

QKI-deleted human bronchial epithelial cells were generated from BEAS-2B cells using the CRISPR/Cas9 system. Further methodological details, including protocols for the clonogenic assay, Annexin V/propidium iodide (PI) staining, mitochondrial reactive oxygen species, mitochondrial membrane potential, metabolic profiling, and genetic rescue experiments using QKI expression plasmids, are described in the online supplement.

### Animal studies

#### Sex as a biological variable

For mouse experiments, both male and female mice were included, and sex distribution did not differ between groups (Fisher’s exact test). For FlexiVent-based pulmonary function testing, only female mice were analyzed in both groups to minimize confounding from sex-associated differences in body weight at the same age, which can influence measured lung mechanics (e.g., compliance) (51). For quantitative analyses of TOMM20 and ATP5A immunostaining, only female mice were analyzed in both groups to minimize sex-related confounding, as mitochondrial phenotypes can differ by sex and are influenced by sex steroid hormones (52); therefore, sex-specific effects could not be evaluated for these assays.

### Transgenic Mice

All transgenic mice were maintained on a C57BL/6 genetic background. To evaluate the effect of QKI deletion specifically in lung epithelial cells, we generated *Sftpc-Cre^/tm1(cre)Ifo^/Qki^flox/flox^* mice (here termed QKI^Δ/Δ^) and *Qki^flox/flox^* mice (here termed control).

To evaluate the effects of QKI deletion specifically in AT2 cells after maturation, we generated *Sftpc^tm1(cre/ERT2)Blh^*/*ROSA26*-mTmG mice (here termed iQKI^wt/wt^) and triple-transgenic *Sftpc^tm1(cre/ERT2)Blh^*/*ROSA26*-mTmG/Qki*^flox/flox^*mice (here termed inducible QKI knockout: iQKI^Δ/Δ^). Further details including mating strategy and experiment outlines are described in the online supplement.

### Pathological lung tissue analysis and lung function measurement

For morphologic analysis of alveoli, the left lung was inflation-fixed at 25 cmH_2_O with 10% neutral buffered formalin and embedded in paraffin.

Airspace enlargement was assessed semi-automatically in QKI^Δ/Δ^ mice aged four weeks and ten to twelve weeks, as well as in iQKI^Δ/Δ^ mice ten weeks after tamoxifen administration. The evaluation used chord length calculated by DeepMasker (53) with H&E-stained lung tissue sections. In addition, immunostaining to assess QKI expression and mitochondrial abundance in alveolar epithelial cells was performed using the same methods as those used for human samples. The number of AT2 cells and the thickness of alveolar walls in newborn (day 0) mice were evaluated using immunostaining and H&E-stained sections with ImageJ. Mouse pulmonary function was measured using the FlexiVent system (SCIREQ, Montreal, QC, Canada). Further details are described in the online supplement.

### Alveolar type 2 epithelial cell isolation, mRNA sequencing, proteome analysis, mtDNA quantification, and spheroid assay

Murine AT2 cell isolation was performed as previously described (54). Isolated AT2 cells were submitted to the Pitt Proteomics Core for proteomic analysis and to Novogene (Sacramento, CA, USA) for mRNA sequencing (RNA-seq). RNA-seq and proteomic analyses were performed using independent control and QKI^Δ/Δ^ AT2 cell samples for each omics dataset (control, n = 4; QK^IΔ/Δ^, n = 4 per dataset). A schematic overview of the multi-omics analysis workflow is shown in Fig. 4A, and additional analytical details are described in the online supplement.

Mitochondrial DNA (mtDNA) content in isolated AT2 cells was quantified using qPCR as described previously (55). To evaluate the characteristics of QKI^Δ/Δ^ AT2 cells, isolated AT2 cells were subjected to three-dimensional *ex vivo* culture to assess the colony formation efficiency (CFE), cell proliferation, and apoptosis. Further details are described in the online supplement.

### RNA-seq analysis

Primary AT2 cells isolated from QKI^Δ/Δ^ and control mice were subjected to RNA-seq analysis. Differential gene expression analysis was performed in R (version 4.6.0) using DESeq2. Differential expression results were summarized as log_2_ fold change, Wald test P value, and Benjamini–Hochberg adjusted P value (FDR). Principal component analysis (PCA) was performed using variance-stabilized expression values. Gene set enrichment analysis (GSEA) was performed using Hallmark gene sets (MSigDB; further details are provided in the online supplement).

### Proteomic analysis

Proteomic analysis was performed using isolated AT2 cells from independent biological cohorts of QKI^Δ/Δ^ and control mice. Differential protein abundance analysis was performed using normalized protein intensity values. Protein abundance changes were summarized as log2 fold change, raw P value, and Benjamini–Hochberg adjusted P value (FDR). PCA was performed using log2-transformed normalized protein intensities *(further details are shown in online supplement)*.

### Integrated transcriptomic–proteomic analysis

RNA-seq and proteomic datasets were integrated at the pathway and gene-symbol levels. Hallmark and GO-BP normalized enrichment scores (NES) derived from RNA-seq and proteomic GSEA were compared to assess transcriptome–proteome concordance. Mitochondrial genes were defined using the MitoCarta3.0 mouse gene inventory. RNA–protein relationships were evaluated by comparing RNA and protein log2 fold changes for shared mitochondrial genes detected in both datasets *(further details are shown in online supplement)*.

### MitoCarta-based mitochondrial gene analysis

Mitochondrial genes were defined using the MitoCarta3.0 mouse gene inventory. A total of 386 mitochondrial genes were identified within the matched RNA–protein dataset. RNA–protein relationships were evaluated by plotting RNA log_2_ fold changes against corresponding protein log_2_ fold changes and calculating Pearson correlation coefficients (further details are provided in the online supplement).

### Statistics

All statistical analyses were performed using JMP 14 software (SAS Institute Inc., Cary, NC, USA) and R (version 4.6.0). Data are presented as mean ± standard error (SE), unless otherwise indicated. P < 0.05 was considered statistically significant for non-omics analyses. For transcriptomic, proteomic, and gene set enrichment analyses, Benjamini–Hochberg FDR < 0.05 was considered statistically significant unless otherwise specified. Additional bioinformatics and statistical details are described in the online supplement and figure legends.

### Study approval

This study was approved by the Institutional Review Board (Protocol ID: 19110180) and the Institutional Animal Care and Use Committee (Protocol IDs: 21048859, 202400592, and 24034592) at the University of Pittsburgh.

## Conflict-of-interest statement

The authors have declared that no conflict of interest exists.

## Author contributions

K.U.: designing the study, acquiring and analyzing data, and writing the draft. K.T., A.M. designing the study, acquiring, analyzing data and interpreting data. L.Z., R.N., and H.U.: acquiring and analyzing data in murine experiments.

K.H.: designing study, and interpreting data, and technical support of experiments.

R.C.K., D.C., P.S., F.S., J.K.A., J.H., C.D.C., and B.A.K.: designing the study, acquiring, analyzing and interpreting data, and making intellectual contributions. N.T.: designing the overall project and specific experiments, analyzing and interpreting data, and editing the draft.

All authors have approved the final version of this manuscript, and none of the authors have any conflicts of interest to declare regarding the publication of this manuscript.

## Funding support

This manuscript was supported, in part, by the US Department of Veterans Affairs, Veterans Health Administration, Office of Research and Development, Biomedical Laboratory Research, and Development by a Merit Review award BX006096-01A1 (T.N.). This work was also supported by NIH R01HL149719 (T.N.) and American Heart Association, Postdoctoral Fellowship program, Grant number 24POST1193659 (K.U.).

## Acknowledgments

The authors are grateful to Li. X, Lakshmi, S.P., and Jiang. S. for data collection and technical assistance.

The authors acknowledge the Unified Flow Cytometry Core at the University of Pittsburgh School of Medicine for providing access to flow cytometry and cell sorting services.

The authors also thank the Health Sciences Mass Spectrometry Core for proteomic analysis support, in part, by the University of Pittsburgh and the Office of the Senior Vice Chancellor for Health Sciences.

K.U. is grateful for the support from the American Heart Association Postdoctoral Fellowship.

## Notes

### Competing Interest Statement

The authors have declared no competing interest.

