## Supplementary Data for "Epithelial QKI Protects Against Emphysema by Maintaining Mitochondrial Integrity"

**Online Supplement**

### Supplemental Methods

#### I. Human studies

i. Analysis of clinical information and *QKI* gene expression in lungs

ii. Immunofluorescence staining and fluorescence intensity for QKI in alveolar type 2 (AT2) epithelial cells

iii. Generation of QKI-deficient BEAS-2B cells

iv. Assessment of colony formation efficiency, cell viability, and apoptosis in QKI-deficient BEAS-2B cells

v. Assessment of metabolic profiles, mitochondrial reactive oxygen species levels, and mitochondrial membrane potential in QKI-deficient BEAS-2B cells

#### II. Animal studies

i. Transgenic mice

ii. Pathological lung tissue image analysis and lung function measurement

iii. Bronchoalveolar lavage fluid (BALF) analysis

iv. AT2 cell isolation, mRNA sequencing, proteomic analysis, mitochondrial DNA abundance measurement, and spheroid assay

v. RNA-seq processing and differential expression analysis

vi. Proteomic data processing and differential abundance analysis

vii. Gene set enrichment analysis

viii. Transcriptome–proteome pathway integration

ix. MitoCarta3.0 gene-level RNA–protein decoupling analysis

x. Source data and software

xi. Immunoblotting in murine AT2 and BEAS-2B cells

**Supplemental Tables**

- Table E1. Reagents used in the present study
- Table E2. Clinical characteristics and sample information for QKI mRNA expression analysis in human lungs
- Table E3. Clinical characteristics and sample information for QKI expression analysis in human AT2 cells
- Table E4. Inflammatory cell numbers in bronchoalveolar lavage fluid

Supplemental Figure Legends

- Figure E1. Timeline of mouse experiments
- Figure E2. Representative immunofluorescence images of human lung sections
- Figure E3. Representative immunofluorescence images of lung sections from control and QKIΔ/Δ mice

- Figure E4. Validation of QKI knockout and add-back in BEAS-2B cells
- Figure E5. Quality control and GO-BP sensitivity analyses for transcriptome–proteome integration
- Figure E6. Myeloid cell–specific QKI deletion does not increase airspace size or alter pulmonary mechanics

### I. Human studies

The lung tissue microarray data with clinical data was obtained from Lung Tissue Research Consortium (LTRC). The FFPE lung tissue sections with clinical data for our microscopic study were obtained from NIH.

#### Analysis of clinical information and *QKI* gene expression in human lungs

We obtained a total of 327 patients' *QKI* gene expression dataset. 72 data were excluded due to post bronchodilator lung function data deficiency, then we analyzed 255 patients' dataset.

#### Immunofluorescence staining and fluorescence intensity for QKI in alveolar type 2 epithelial (AT2) cells

The lung tissues were obtained as FFPE sections and stained based on previously described (1). Images were acquired using a Nikon Ti microscope (Nikon Instruments Inc., Melville, NY, USA) under the same conditions across all batches. AT2 cells were identified as DC-LAMP-positive cells (2). Regions of interest (ROIs) were defined by encompassing the entire DC-LAMP-positive cells, and cells without DAPI nuclear staining were excluded from the analysis. The areas and QKI fluorescence intensities were measured using ImageJ

(National Institutes of Health, Bethesda, MD, USA).

At least fifty AT2 cells were evaluated from at least three different fields of view in each sample. The average QKI expression for each sample was calculated. To account for batch effects, QKI expressions were normalized by dividing each sample's QKI expression by the average QKI expression of non-COPD samples within the same batch and calculated as relative QKI expression in AT2.

***In vitro* experiments; QKI deleted Beas 2B cells ($QKI^{-/-}$ B2B cells)**

BEAS-2B cells were purchased from American Type Culture Collection (Manassas, VA, USA) and maintained in Keratinocyte SFM (Thermo Fisher Scientific, Waltham, MA, USA).

We generated CRISPR/Cas9 QKI knockout BEAS-2B cell lines with the following steps. We designed guide RNAs for the exon 2 of QKI using CRISPOR (https://crispor.gi.ucsc.edu/)(3) and purchased oligos from Integrated DNA Technologies, Inc. (Coralville, IA, USA) (Forward: 5'-CACCGACAATAGGTCCCACAGCATC-3', Reverse: 5'-AAACGATGCTGTGGGACCTATTGTC-3'). We used the lentiGuide-Puro (Addgene, plasmid #52963, Watertown, MA, USA) for the lentiviral vector. According to the Zhang lab protocol (4, 5), we performed ligation of the oligos into the vector plasmids and transformed the ligated plasmids into Stbl3 E. coli (Invitrogen). After incubation, we purified the vector plasmids and verified the sequence.

In lentiviral vector packaging, we purchased Lenti-X 293T (Takara Bio USA, Inc., Mountain View, CA, USA) cells and maintained in DMEM (Thermo Fisher Scientific) containing non-essential amino acids, sodium pyruvate, penicillin/streptomycin, and 10% fetal bovine serum. We transfected the vector plasmids, psPAX2 (Addgene, plasmid #12260), and pMD2.G (Addgene, plasmid #12259) into Lenti-X 293T cells using Lipofectamine 3000 (Thermo Fisher Scientific). After incubation for 48 hours, we collected the lentiviral vector.

For transduction of the lentiviral vector, we used Cas9-expressing BEAS-2B cells kindly provided by Alder lab. After 48 hours from transduction, we started the selection of the CRISPR/Cas9 QKI knockout BEAS-2B cells using puromycin. After the selection, we performed limiting dilution cloning to obtain single cell colonies and confirmed the deletion of QKI expression by immunoblotting.

In the QKI restoration experiments, we generated a QKI-expressing lentiviral vector using QKI in pLX304 (DNASU, clone ID: HsCD00441520, Tempe, AZ, USA) psPAX2, and pMD2.G as shown above. Then, we transduced into the CRISPR/Cas9 QKI knockout BEAS-2B cells (QKI add-back B2B) and obtained single cell colonies.

**Assessments of colony formation efficiency, cell viability, and apoptosis**

Colony formation efficiency was assessed as plating efficiency (PE) in each cell type were measured by a clonogenic assay (6); Cells were seeded at a density of 200 cells per well in a 6-well plate and cultured for 10 days. Cells were stained with crystal violet, and the number of colonies was counted. A colony was defined as a cluster consisting of ≥50 cells. Plating efficiency (%) was calculated as the ratio of the number of colonies formed to the initial number of seeded cells. Each experimental batch included two wells per group, and the mean value was determined. The assay was performed in three independent experiments, and statistical analyses were conducted accordingly.

Cell viability was assessed by MTT assay as previously described (7); Briefly, after overnight incubation following cell seeding ($1.2 \times 10^5$ cells), cells were stained with 3-(4,5-dimethylthiazol-2-yl)-2,5-diphenyltetrazolium bromide (Sigma-Aldrich, St. Louis, MO, USA) and the absorbance was measured at 560 nm – 750 nm using a Synergy H1 Microplate Reader（Agilent Technologies, Inc., Winooski, VT, USA）. The absorbance of $QKI^{-/-}$ B2B and QKI add-back B2B cells were evaluated relative to the control, which was set to 100% in each experimental batch. Each experimental batch included three wells per group, and the mean value was determined. The assay was performed in three independent experiments, and statistical analyses were conducted accordingly. Cell apoptosis was evaluated using an Annexin V-PI staining kit (Biolegend, San Diego, CA, USA) and analyzed using an LSR Fortessa Cell Analyzer (BD Biosciences, Franklin Lakes, NJ, USA). The fraction was regarded as below; AnnexinV-PI- cells: live cells, AnnexinV+PI- cells: early apoptotic cells, Annexin+PI+ cells: late apoptotic cells, Annexin+PI+ cells: dead cells. The assay was performed in three independent experiments, and statistical analyses were conducted accordingly.

**Assessments of mitochondrial reactive oxygen species levels, mitochondrial membrane potential, and metabolic profiles**

The metabolic status of $QKI^{-/-}$ B2B cells was evaluated using the Seahorse XFe96 Analyzer (Agilent). Cells were seeded at a density of $6.0 \times 10^4$ cells per well and incubated overnight before the assay. Oxygen consumption rate (OCR) and extracellular acidification rate (ECAR) were assessed to determine mitochondrial and glycolytic activity, respectively.

For mitochondrial stress testing, the following compounds were sequentially injected at specified time points:

- Oligomycin (2 μM): Added after measurement 3 to inhibit ATP synthase.
- FCCP (1 μM): Added after measurement 6 to uncouple mitochondrial oxidative phosphorylation.
- Rotenone + Antimycin A (0.5 μM each): Added after measurement 9 to inhibit complex I and III, shutting down mitochondrial respiration.

OCR and ECAR values were normalized to cell number using a DNA-binding fluorescence assay, which provides an estimate of cell number based on DNA content. Eight wells per group were analyzed, and outliers were excluded from the final dataset.

The levels of mitochondrial reactive oxygen species (ROS) and membrane potential were measured by Mitochondrial Superoxide Assay Kit (ab219943, Abcam, Cambridge, MA, USA) and Mitochondrial Membrane Potential Assay Kit (ab113852, Abcam). Fluorescence intensity of $QKI^{-/-}$ B2B and QKI add-back B2B cells were evaluated relative to the control, which was set to 100% in each experimental batch. For each batch, the mean fluorescence intensity was calculated from six wells per condition. The experiment was repeated three times independently, and statistical analyses were performed.

### II. Animal studies

The Institutional Animal Care and Use Committee approved the study (ID: 21048859 and 202400592).

#### Transgenic Mice

All transgenic mice were maintained in a C57/BL6 genetic background.

Double-transgenic *Sftpc-Cre*$^{/tm1(cre)}$/*Qki*$^{flox/flox}$ mice were generated from *Qki*$^{flox/flox}$ (kindly gifted by Dr. Hu) and *Sftpc-Cre*$^{/tm1(cre)}$ mice (kindly gifted by Dr. Alder (8)). The double-transgenic mice were bred with *Qki*$^{flox/flox}$ mice to generate *Sftpc-Cre*$^{+/tm1(cre)}$/*Qki*$^{flox/flox}$ mice (here termed $QKI^{\Delta/\Delta}$) and *Qki*$^{flox/flox}$ mice (here termed control). In the $QKI^{\Delta/\Delta}$ mice, QKI is eliminated in alveolar and bronchial epithelial cells during whole period.

Also, to evaluate the effect of QKI in myeloid cells including alveolar and interstitial macrophages, we also generate myeloid cell specific QKI deletion mice by breeding *Qki*$^{flox/flox}$ mice with *Lyz2*$^{tm1(cre)Ifo}$/J mice (Stock No: 004781) purchased from The Jackson Laboratory (Bar Harbor, ME, USA) to generate *Lyz2*$^{+/tm1(cre)Ifo}$ /*Qki*$^{flox/flox}$ mice (here termed Lyz2-QKI$^{\Delta/\Delta}$) and *Qki*$^{flox/flox}$ mice (here termed Lyz2-control) same as the *Sftpc-Cre*$^{+/tm1(cre)}$/*Qki*$^{flox/flox}$ mice breeding strategy.

To evaluate lung epithelial cell differentiation and investigate the effects of QKI deletion specifically in alveolar type 2 epithelial cells (AT2 cells) after maturation, *Sftpc*$^{tm1(cre/ERT2)Blh}$/*ROSA26*-mTmg mice were bred with *Qki*$^{flox/flox}$ mice to generate double-transgenic *Sftpc*$^{tm1(cre/ERT2)Blh}$/*ROSA26*-mTmg mice (kindly gifted by Alder lab (8)) (here termed iQKI$^{wt/wt}$) and triple-transgenic *Sftpc*$^{tm1(cre/ERT2)Blh}$/*ROSA26*-mTmg/*Qki*$^{flox/flox}$ mice (here termed inducible QKI knockout: iQKI$^{\Delta/\Delta}$). These mice were administered tamoxifen (Thermo Fisher Scientific, 200 mg/kg) three times to induce Cre-mediated recombination at eight to nine weeks of age. This approach allowed us to specifically delete QKI in AT2 cells after lung development had reached maturity, enabling the assessment of QKI's role in maintaining alveolar integrity and its contribution to emphysema pathogenesis.

**Pathological lung tissue image analysis and lung function measurement**

For morphologic analysis in alveoli, mice left lungs were inflation fixed at 25 cm $H_2O$ with 10% neutral buffered formalin and were embedded in paraffin. Tissue sections (4-μm thick) were immunostained with antibodies (see Table E1 for a list of antibodies) as previously described (1), and fluorescence images were obtained using a Nikon Ti confocal microscope (Nikon).

Airspace enlargement was assessed semi-automatically in QKI$^{\Delta/\Delta}$ mice aged four weeks and 10 to 12 weeks, as well as in iQKI$^{\Delta/\Delta}$ mice 10 weeks after tamoxifen treatment. The evaluation utilized the chord length calculated by DeepMasker (9) with H&E-stained lung tissue sections.

For mitochondrial quantitative analysis, we measured the fluorescence intensities of translocase of outer mitochondrial membrane 20 (TOMM20) and ATP synthase F1 subunit alpha (ATP5A). A minimum of ten AT2 cells were measured per sample for evaluation. Images were obtained using a Nikon Ti confocal microscope (Nikon), and the mean gray value for each channel was calculated within the ROIs corresponding to individual DC-LAMP positive AT2 cells using ImageJ software (National Institutes of Health), and the average value per sample was used for statistical comparisons.

To evaluate the influence of QKI deficiency on embryonic lung maturation, we collected thoracic cavities from newborn (day 0) $QKI^{\Delta/\Delta}$ and control mice, fixed them by inflation with 10% neutral buffered formalin, and embedded the tissues in paraffin. We then prepared 4-μm sections and immunostained them with antibodies. The number of AT2 cells per alveolar perimeter length was quantified, while excluding any areas containing airways, blood vessels, or other non-alveolar structures to ensure specificity. In addition, to assess lung structural maturation, we measured alveolar wall thickness in fifteen randomly selected alveolar walls per sample (10-12). Specifically, any regions in which the alveolar wall appeared double-layered, indistinct, or contained non-alveolar tissues were omitted to maintain accuracy, and each alveolar wall was measured at an approximately right angle to avoid artifactual thickening from oblique sectioning.

Pulmonary function in mouse lung was measured using Flexivent system (SCIREQ, Montreal, QC, Canada) (13). All animal procedures were performed according to a protocol approved by the University of Pittsburgh IACUC.

**Bronchoalveolar lavage fluid (BALF) analysis**

BALF was obtained using 2×0.8 ml (1.6 ml in total) aliquots of PBS. After a cell count was performed, cell fractions in BALF samples were analyzed by flow cytometry (FCM). The details were previously described (1, 14) (also see Table E2 for a list of antibodies and fluorochromes).

Data were acquired using an LSR Fortessa Cell Analyzer (BD Biosciences) and were analyzed by FlowJo software (TreeStar, Ashland, OR, USA).

$CD45^+CD11b^-SiglecF^+$ cells were regarded as alveolar macrophages,

CD45$^{+}$CD11b$^{+}$Ly6G$^{+}$ cells were regarded as neutrophils, CD45$^{+}$CD11b$^{+}$Ly6G$^{-}$SiglecF$^{-}$ cells were regarded as interstitial macrophages, and CD45$^{+}$CD11b$^{-}$Ly6G$^{-}$SiglecF$^{+}$CD11c$^{-}$ cells were regarded as eosinophils. In the remaining CD45$^{+}$ fractions, CD19$^{+}$CD3ε$^{-}$ cells were regarded as B cells and CD19$^{+}$CD3ε$^{+}$ cells were regarded as T cells.

**AT2 cell isolation, mRNA sequencing, proteome analysis, the measurement of abundance of mitochondrial DNA, and spheroid assay**

Murine AT2 cell isolation was performed as previously described (15); briefly, mice lungs were individualized using dispase (Corning, NY, USA) and gentleMACS$^{TM}$ Dissociator (Miltenyl Biotec, Bergisch Gladbach, NRW, Federal Republic of Germany), and stained with CD45, PECAM, EpCAM, and MHC class2 (BD Biosciences). CD45$^{-}$PECAM$^{-}$EpCAM$^{+}$MHC class2$^{+}$ cells were regarded as AT2 cells and sorted using the BD Aria II at the Unified Flow Cytometry Core, University of Pittsburgh School of Medicine. Total RNA was isolated using a RNeasy plus mini kit (QIAGEN, Valencia, CA), and mRNA sequencing was performed by NOVO-GENE (Sacramento, CA, USA), the sequencing data was provided from NOVOmagic. The proteomics analysis for sorted AT2 cells were performed in the Health Sciences Mass Spectrometry Core (RRID:SCR_025222), and the sequencing data was analyzed using DAVID. Isolated AT2 cells were also used for spheroid assay as the same method as previously described (16) (17); One thousand AT2 cells were seeded in Matrigel (Corning) and cultivated for ten days, and colony formation efficiency was calculated (The number of spheres (>50 um)/1,000 x 100). To evaluate cell proliferation and cell apoptosis in mice AT2 cells, AT2 cells obtained from spheroids were stained with Ki-67 (Biolegend) and Annexin V-PI staining (Biolegend) and analyzed using an LSR Fortessa Cell Analyzer (BD Biosciences). The fraction was regarded as below; AnnexinV$^{-}$PI$^{-}$ cells: live cells, AnnexinV$^{+}$PI$^{-}$ cells: early apoptotic cells, Annexin$^{+}$PI$^{+}$ cells: late apoptotic cells, Annexin$^{+}$PI$^{+}$ cells: dead cells.

**RNA-seq processing and differential expression analysis**

mRNA sequencing (RNA-seq) was performed using AT2 cells isolated from independent control and $QKI^{\Delta/\Delta}$ mice. Control and $QKI^{\Delta/\Delta}$ RNA-seq samples were generated independently from the proteomic samples, with 4 biological replicates per group. RNA-seq libraries were prepared by Novogene using a poly(A)-enriched directional mRNA library preparation. Library strandedness was confirmed from BAM files using RSeQC infer_experiment.py (18), and gene-level counts were generated using featureCounts with paired-end reverse-stranded counting against the mouse GRCm39 annotation (19).

Differential gene expression was analyzed in R using DESeq2 (20), with control samples used as the reference group. Genes with very low total counts were removed before differential expression modeling. Differential expression results were reported as log2 fold change, Wald test P value, and Benjamini–Hochberg adjusted P value, hereafter referred to as FDR. Throughout these analyses, log2FC refers to the log2 fold change for $QKI^{\Delta/\Delta}$ versus control. RNA-seq volcano plots were generated using all annotated genes with available log2 fold change and adjusted P value. Genes were highlighted as upregulated or downregulated in $QKI^{\Delta/\Delta}$ when Benjamini–Hochberg FDR $< 0.05$ and $|log2FC| \geq 1$.

RNA-seq principal component analysis (PCA) was performed using variance-stabilized expression values from expressed genes. PCA was performed using prcomp in R.

**Proteomic data processing and differential abundance analysis**

Proteomic analysis was performed by the Pitt Proteomics Core using AT2 cells isolated from independent control and $QKI^{\Delta/\Delta}$ mice, with 4 biological replicates per group. Proteomic samples were generated independently from the RNA-seq samples. Proteomic results were analyzed at the protein group level using the LeadingRazorProtein annotation provided in the proteomics output. For gene set enrichment and RNA–proteome integration, protein groups were mapped to gene symbols. Differential protein abundance analysis was performed using normalized protein intensity values. Protein abundance changes were summarized as log2 fold change for $QKI^{\Delta/\Delta}$ versus control, raw P value, and Benjamini–Hochberg FDR.

For MA plot visualization, A was calculated as the mean log2(normalized intensity + 1) across all proteomic samples, and M was calculated as the difference between the mean log2(normalized intensity + 1) in $QKI^{\Delta/\Delta}$ and control samples. Proteins with Benjamini–Hochberg FDR < 0.05 were highlighted.

Proteome PCA was performed using log2-transformed normalized protein intensities. To minimize the influence of zero or missing values, PCA was performed using protein groups quantified in all samples and showing nonzero variance after log2 transformation. This retained 2,043 protein groups for PCA. PCA was performed using prcomp in R.

**Gene set enrichment analysis (GSEA)**

Preranked gene set enrichment analysis was performed in R using clusterProfiler (21). MSigDB Hallmark gene sets were used for the main pathway analysis, and Gene Ontology Biological Process gene sets were used as a sensitivity analysis. RNA-seq GSEA was performed using a protein-coding ranked list to improve comparability with the proteomic dataset. An all-gene RNA ranked list was retained in the supplementary source data for transparency but was not used for the primary GSEA figures. Proteomic GSEA was performed using representative gene symbols assigned to protein groups.

For both RNA-seq and proteomic GSEA, genes or proteins were ranked using the following signed significance metric:

$$\text{sign}(\log_2 \text{ fold change}) \times -\log_{10}(\text{raw } P \text{ value})$$

GSEA results were summarized using normalized enrichment score, nominal P value, Benjamini–Hochberg adjusted P value, and core enriched genes. Terms with $FDR < 0.05$ were considered significant unless otherwise specified. For GO Biological Process plots, significant GO terms were further filtered using simplify in clusterProfiler, with a semantic similarity cutoff of 0.7, to reduce redundancy before visualization. Full GO-BP GSEA results were retained in the supporting data.

**Transcriptome–proteome pathway integration**

For pathway-level transcriptome–proteome integration, RNA-seq and proteomic GSEA results were matched by gene set name for Hallmark analysis and by GO term ID for GO Biological Process analysis (22). Normalized enrichment scores from RNA-seq and proteome analyses were plotted against each other. Because RNA-seq and proteomic measurements were obtained from independent sample sets, integration was performed at the gene symbol or pathway level rather than as a sample-paired analysis.

In the Hallmark NES comparison, selected mitochondrial/metabolic Hallmark pathways were annotated for visualization. Hallmark mitochondrial/metabolism-related terms were defined conservatively as oxidative phosphorylation, fatty acid metabolism, and reactive oxygen species pathway. In the GO Biological Process NES comparison, mito/metabolism-related terms were annotated using keyword matching, including mitochondrial, respiratory chain, oxidative phosphorylation, aerobic respiration, tricarboxylic acid cycle, fatty acid oxidation, lipid oxidation, organic acid catabolism, carboxylic acid catabolism, monocarboxylic acid catabolism, acyl-CoA, and CoA metabolism.

**MitoCarta3.0 gene-level RNA–protein decoupling analysis**

Mitochondrial genes were defined using the mouse MitoCarta3.0 gene inventory (23). RNA-seq and proteomic differential results were matched by gene symbol after removal of entries without gene symbols or log2 fold change values. When duplicate gene symbols were present within either omics dataset, the entry with the largest absolute log2 fold change was retained before RNA–proteome matching. This yielded 2,225 shared gene symbols between the RNA-seq and proteomic datasets.

MitoCarta3.0 genes within the shared RNA–proteome gene set were then extracted for gene-level mitochondrial RNA–protein comparison. For each gene, RNA–protein discrepancy was calculated as: RNA log2FC − protein log2FC.

The final MitoCarta3.0-matched dataset contained 386 genes. Median RNA log2FC, median protein log2FC, and median RNA–protein discrepancy were calculated across these genes. Deviation of the RNA–protein discrepancy from zero was assessed using a two-sided Wilcoxon signed-rank test.

**Source data and software**

Source data for Figure 4 and Figure E5 were provided as Excel files. Underlying values for each figure panel were organized in separate worksheets, with variable definitions included in a metadata worksheet. Full RNA-seq differential expression results and full proteomic differential abundance results were provided as a separate supplementary table.

Analyses were performed using R version 4.6.0. R packages used for the omics analyses and visualization included DESeq2, clusterProfiler, msigdbr, dplyr, ggplot2, ggrepel, and writexl.

**Immunoblotting in murine AT2 and Beas 2B cells**

To measure protein expression levels, isolated mice AT2 and Beas 2B cells were lysed directly by RIPA buffer (Sigma-Aldrich) with protease and phosphor inhibitor (Thermo Fisher Scientific) and extracted proteins (24). In mice study, protein levels were assessed relative to the level of β-actin using ImageJ software (National Institutes of Health), and performed statistical analysis.

**Table E1: Reagents list for the present study**

**Antibodies for immunostaining and immunoblotting**

| Antigen | Cat. no. | Company | Titer (for IF) | Titer (for immunoblotting) |
|---|---|---|---|---|
| QKI | A300-183A | Bethyl Laboratories, (Montgomery, TX, USA) | 1:500<br>1:1,000 | 1:5,000 |
| Podoplanin | ab11936 | Abcam | 1:100 | N/A |
| DC-LAMP | DDX0191P | Novus Biologicals, (Centennial, CO, USA) | 1:100 | N/A |
| CC10 (Scgb1a1) | sc-390313 | Santa Cruz (Dallas, TX, USA) | 1:1,000 | N/A |
| Acetylated tubulin | T7451 | Sigma-Aldrich | 1:1,000 | N/A |
| GFP | ab13970 | Abcam | 1:1,000 | N/A |
| F4/80 | 14-4801-82 | Thermo Fisher Scientific | 1:100 | N/A |
| SP-C | sc-518029 | Santa Cruz | 1:200 | N/A |
| GFP | ab13970 | Abcam | 1:1,000 | N/A |
| TOMM20 | ab186735 | Abcam | 1:200 | N/A |
| ATP5A | ab14748 | Abcam | 1:400 | N/A |
| Cleaved Caspase3 | 9661S | Cell Signaling Technology (Danvers, MA, USA) | N/A | 1:1,000 |
| V5 Tag | R960-25 | Thermo Fisher Scientific | N/A | 1:5,000 |
| β-actin | 60000-Ig | Proteintech Group (Rosemont, IL, USA) | N/A | 1:25,000 |

**Antibodies for flow cytometry**

| Antigen | Cat. no. | Company | Fluorochrome |
|---|---|---|---|
| CD16/32 | 156604 | Biolegend | Purified |
| CD45 | 103108 | Biolegend | FITC |
| SiglecF | 155508 | Biolegend | APC |
| Ly6G | 127612 | Biolegend | Pacific Blue |
| CD11b | 101208 | Biolegend | PE |
| CD19 | 117324 | Biolegend | PerCP-Cy5.5 |
| CD3ε | 100320 | Biolegend | PE-Cyanine 7 |
| Ki-67 | 11-5698-82 | Thermo Fisher Scientific | FITC |

**Antibodies for alveolar type 2 epithelial cell isolation**

| Antigen | Cat. no. | Company | Fluorochrome |
|---|---|---|---|
| PECAM-1 | 25-0311-82 | Thermo Fisher Scientific | PE-Cyanine 7 |
| CD45 | 25-0451-82 | Thermo Fisher Scientific | PE-Cyanine 7 |
| MHC class II | 48-5321-82 | Thermo Fisher Scientific | eFluor 450 |
| EpCAM | 17-5791-82 | Thermo Fisher Scientific | APC |
| 7-AAD | 555816 | BD Biosciences | 7-AAD |

**Table E2: Clinical characteristics of patients and sample information in the analysis of *QKI mRNA* expression in human lungs**

| Items | Non-COPD N = 83 | GOLD I+II N = 106 | GOLD III+IV N = 66 | P value (ANOVA) |
|---|---|---|---|---|
| *QKI* probe intensity | 6.42 ± 0.03 | 6.43 ± 0.03 | 6.23 ± 0.06* | <0.001 |
| Age (year) | 62.5 ± 1.0 | 67.9 ± 0.9* | 61.3 ± 1.2 | <0.001 |
| Female (n) | 46 | 41 | 30 | 0.07 |
| Smoking status (Never/Ever) | 24/48 n = 72 | 5/98 n = 103 | 1/65 n = 66 | <0.0001 |
| Smoking index (pack-years) | 24.5 ± 4.2*, n = 72 | 55.6 ± 3.5, n = 103 | 49.3 ± 4.4, n = 66 | <0.001 |
| GOLD stage Non-COPD/I/II/III/IV (n) | 83/0/0/0/0 | 0/20/86/0/0 | 0/0/0/28/38 | <0.0001 |
| $FEV_1$ (% of predicted) | 99.2 ± 1.3 | 69.1 ± 1.1 | 29.8 ± 1.4 | <0.0001[†] |
| FVC (% of predicted) | 96.8 ± 1.3 | 89.1 ± 1.3 | 66.4 ± 2.0 | <0.0001[†] |

*Definition of abbreviations:* GOLD = Global Initiative for Chronic Obstructive Lung Disease; COPD = chronic obstructive pulmonary disease; $FEV_1$ = forced expiratory volume in 1 second; FVC = forced vital capacity; N.S. = not significant.

The table shows the clinical characteristics of patients with or without COPD. Pulmonary function test data were collected after bronchodilator treatment. The diagnosis of COPD was confirmed by pulmonologists.

Continuous variables (*QKI* probe intensity, age, smoking index, $FEV_1$, % of predicted $FEV_1$, and % of predicted FVC) are presented as the mean ± SEM and were analyzed using one-way ANOVA followed by pairwise testing with the Tukey-Kramer HSD test for comparisons. Categorical variables were analyzed with chi-square tests.

* indicates that the group is significantly different ($P < 0.05$, Tukey-Kramer HSD test) compared to the other groups.

† indicates that all groups are significantly different ($P < 0.05$, Tukey-Kramer HSD test) from each other.

**Table E3: Clinical characteristics and sample information for the analysis of QKI expression in human alveolar type 2 epithelial cells (AT2 cells)**

| Items | Non-COPD N = 10 | GOLD III+IV N = 10 | P value |
|---|---|---|---|
| Age (year) | 71.6 ± 2.3 | 59.7 ± 2.1 | <0.005 |
| Female (n) | 5 | 7 | 0.35 |
| Smoking index (pack-years) | 52.0 ± 8.4 | 45.6 ± 10.0 | 0.63 |
| Smoking status (Never/Ever, n) | 0/10 | 0/10 | N.S. |
| BMI (kg/m$^2$) | 27.5 ± 2.4 | 25.0 ± 1.2 | 0.37 |
| GOLD stage Non-COPD/I/II/III/IV (n) | 10/0/0/0/0 | 0/0/0/2/8 | <0.0001 |
| $FEV_1$ (% of predicted) | 93.9 ± 3.9 | 21.4 ± 2.2 | <0.0001 |
| FVC (% of predicted) | 93.3 ± 4.0 | 53.5 ± 3.3 | <0.0001 |
| $DL_{CO}$, (% of predicted) | 83.1 ± 4.3, n = 8 | 24.1 ± 2.7, n = 6 | <0.0001 |
| Sample location (Upper/middle/lower lobe, n) | 10/0/0 | 10/0/0 | N.S. |
| Number of assessed AT2 cells (n) | 62.2 ± 3.0 | 65.2 ± 3.9 | 0.55 |

*Definition of abbreviations:* GOLD = Global Initiative for Chronic Obstructive Lung Disease; COPD = chronic obstructive pulmonary disease; $FEV_1$ = forced expiratory volume in 1 second; FVC = forced vital capacity; $DL_{CO}$ = Diffusing capacity for carbon monoxide; BMI = body mass index; N.S. = not significant.

The table shows the clinical characteristics of patients with smoking history with or without severe COPD. The pulmonary function test data was collected after bronchodilator treatment. The diagnosis of COPD was confirmed by pulmonologists.

Continuous variables (Agee, smoking index, BMI, $FEV_1$, % of predicted $FEV_1$, % of predicted FVC, % of predicted $DL_{CO,}$ and number of measured AT2 cells) are presented as the mean ± SEM and were analyzed using unpaired Student's t tests. Categorical variables were analyzed with chi-square tests.

**Table E4: Inflammatory cell numbers in bronchoalveolar lavage fluid (BALF)**

| Variables | Control<br>N = 8 | $QKI^{\Delta/\Delta}$<br>N = 5 | P value |
|---|---|---|---|
| Total cells ($x10^4$/mL) | 5.48 ± 1.09 | 11.29 ± 1.82 | <0.05 |
| Alveolar macrophages ($x10^4$/mL) | 5.21 ± 1.05 | 11.06 ± 1.53 | <0.05 |
| Interstitial macrophages ($x10^3$/mL) | 1.20 ± 0.20 | 0.84 ± 0.11 | 0.15 |
| Neutrophils ($x10^3$/mL) | 0.89 ± 0.42 | 0.98 ± 0.20 | 0.85 |
| Eosinophils (/mL) | 73.88 ± 28.54 | 21.95 ± 16.80 | 0.14 |
| B cells ($x10^3$/mL) | 0.41 ± 0.24 | 0.38 ± 0.09 | 0.91 |
| T cells ($x10^3$/mL) | 0.15 ± 0.05 | 0.14 ± 0.05 | 0.85 |

The BALF cell fraction in 10-week-old control and $QKI^{\Delta/\Delta}$ mice was analyzed by flow cytometry, and each cell number was calculated (n = 5~8 per group). Data are presented as the mean ± SEM and were analyzed using Welch's t-test.

### Supplemental Figure

#### Figure E1

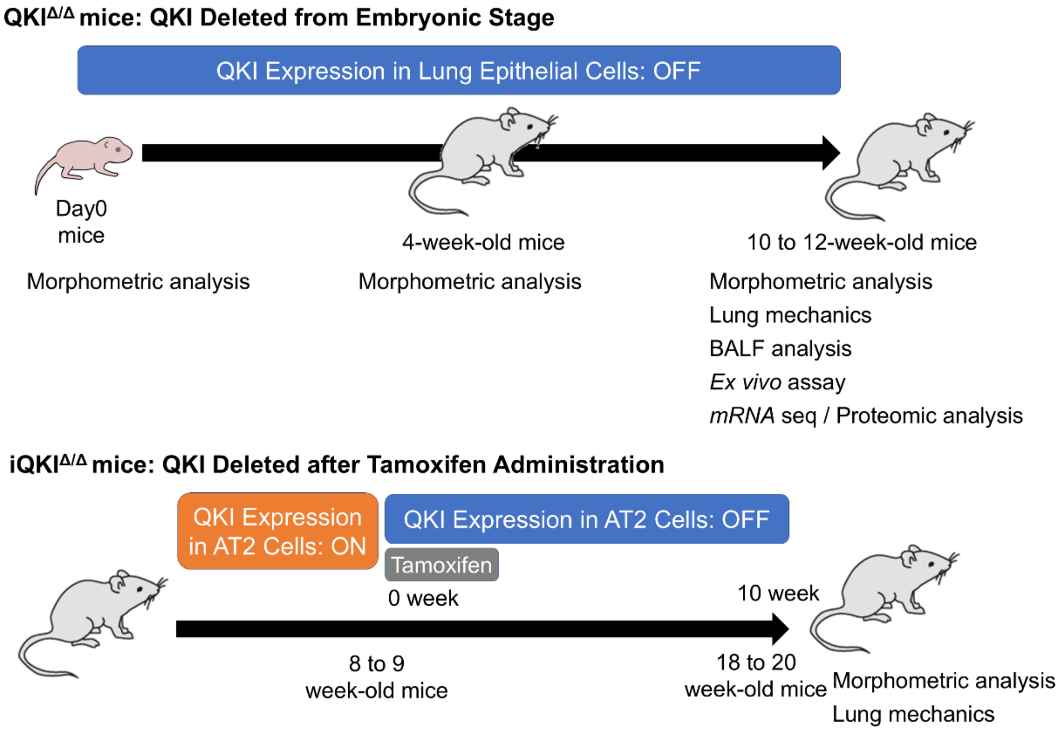


***Timeline of mouse experiments***

The *Sftpc-Cre*$^{+/tm1(cre)}$/*Qki*$^{flox/flox}$: QKI$^{\Delta/\Delta}$ mice, in which QKI was deleted from the embryonic stage, were analyzed at different time points to evaluate the development of emphysema and lung function. Morphometric analysis was performed at postnatal day 0, 4 weeks, and 10 to 12 weeks. At 10 to 12 weeks, additional assessments, including lung mechanics, BALF analysis, *ex vivo* assays, and mRNA-seq/proteomic analysis, were conducted.

For the *Sftpc*$^{tm1(cre/ERT2)Blh}$/*ROSA26*-mTmg/*Qki*$^{flox/flox}$: mTmg: iQKI$^{\Delta/\Delta}$ mice, where QKI deletion was induced after tamoxifen administration, QKI expression remained intact in AT2 cells before tamoxifen treatment. Mice at 8 to 9 weeks of age received tamoxifen, and by 10 weeks, QKI expression was lost in AT2 cells. Morphometric analysis and lung mechanics were evaluated at 18 to 20 weeks.

**Figure E2**

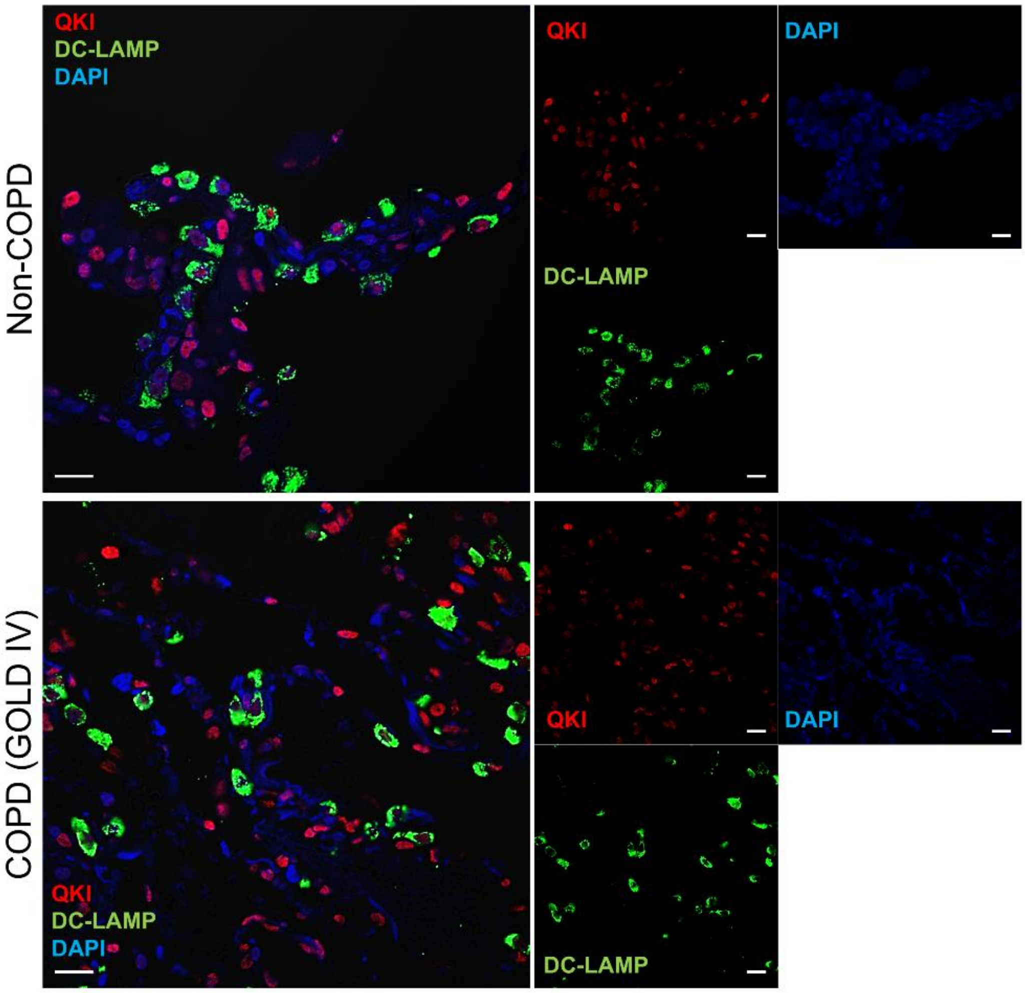


***Representative immunofluorescence images of human lung sections from non-COPD and COPD***

Lung sections were stained for QKI (red), DC-LAMP (green; a marker of alveolar type 2 epithelial cells), and DAPI (blue). Scale bars, 50 um.

**Figure E3**

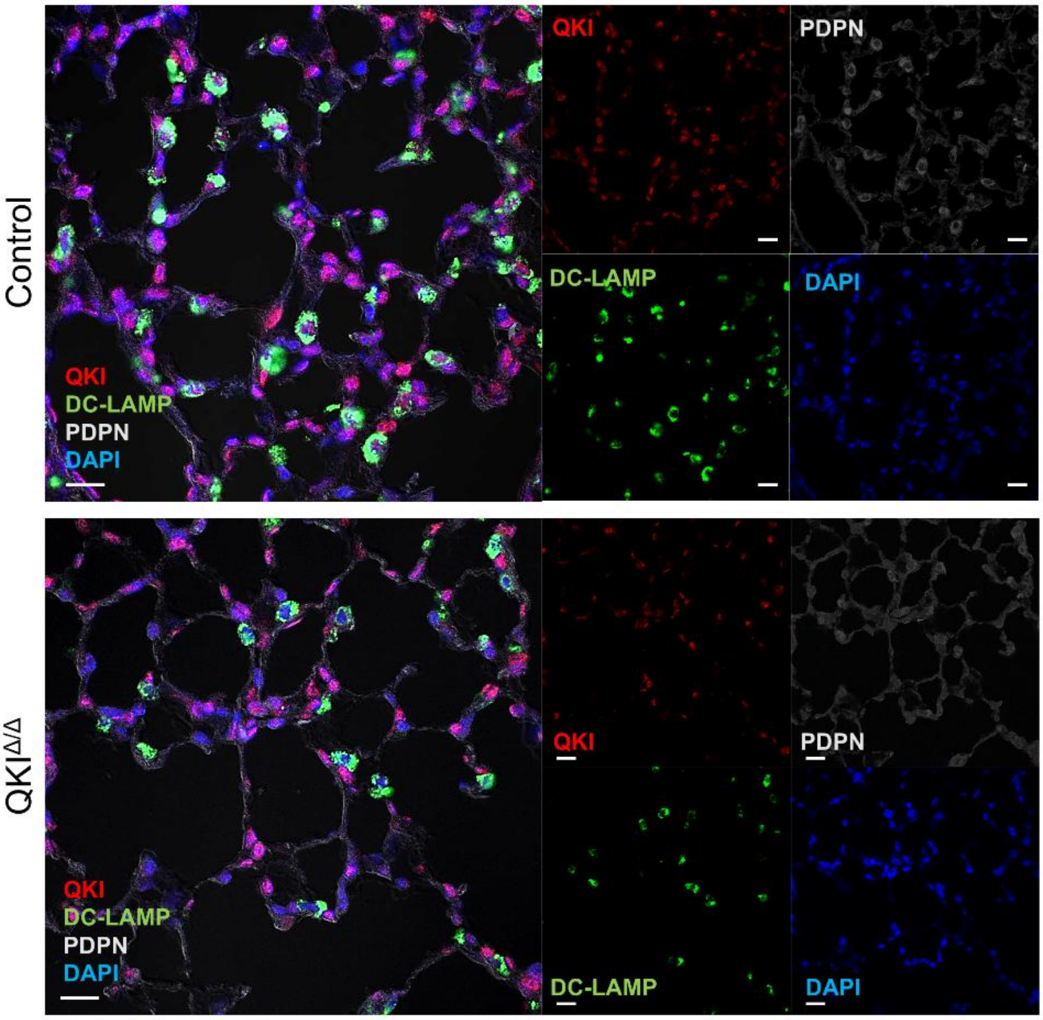


***Representative immunofluorescence images of alveolar regions from control and QKI$^{\Delta/\Delta}$ mice***

Lung sections were stained for QKI (red), DC-LAMP/LAMP3 (green; an alveolar type 2 epithelial cell marker), PDPN/podoplanin (white; an alveolar epithelial cell marker), and DAPI (blue). Scale bars, 50 μm.

**Figure E4**

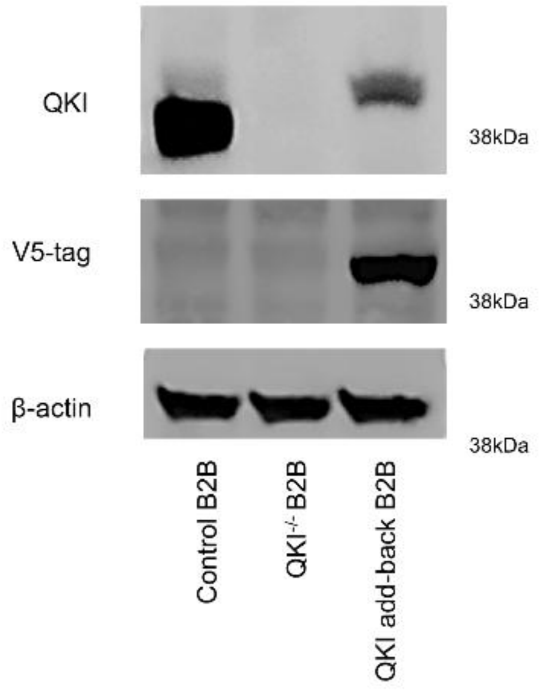


***Validation of QKI knockout and add-back in BEAS-2B cells***

QKI expression: Immunoblotting analysis confirms the complete loss of QKI protein in $QKI^{-/-}$ B2B cells compared to control B2B cells. In QKI add-back B2B cells, QKI expression was restored, but the recovery was incomplete, indicating partial reconstitution by the plasmid-based add-back system.

V5-tag expression: The V5-tag blot demonstrates that QKI protein introduced into QKI add-back B2B cells was ectopically expressed, as evidenced by the presence of the V5-tag signal. This finding confirms that the detected QKI protein in the add-back condition originated from the plasmid-based construct rather than endogenous expression. The QKI band shift in the QKI add-back B2B lane suggests that the V5 tag increased the apparent molecular weight of recombinant QKI protein, leading to a slightly different migration pattern compared with endogenous QKI in control B2B cells.

**Figure E5**

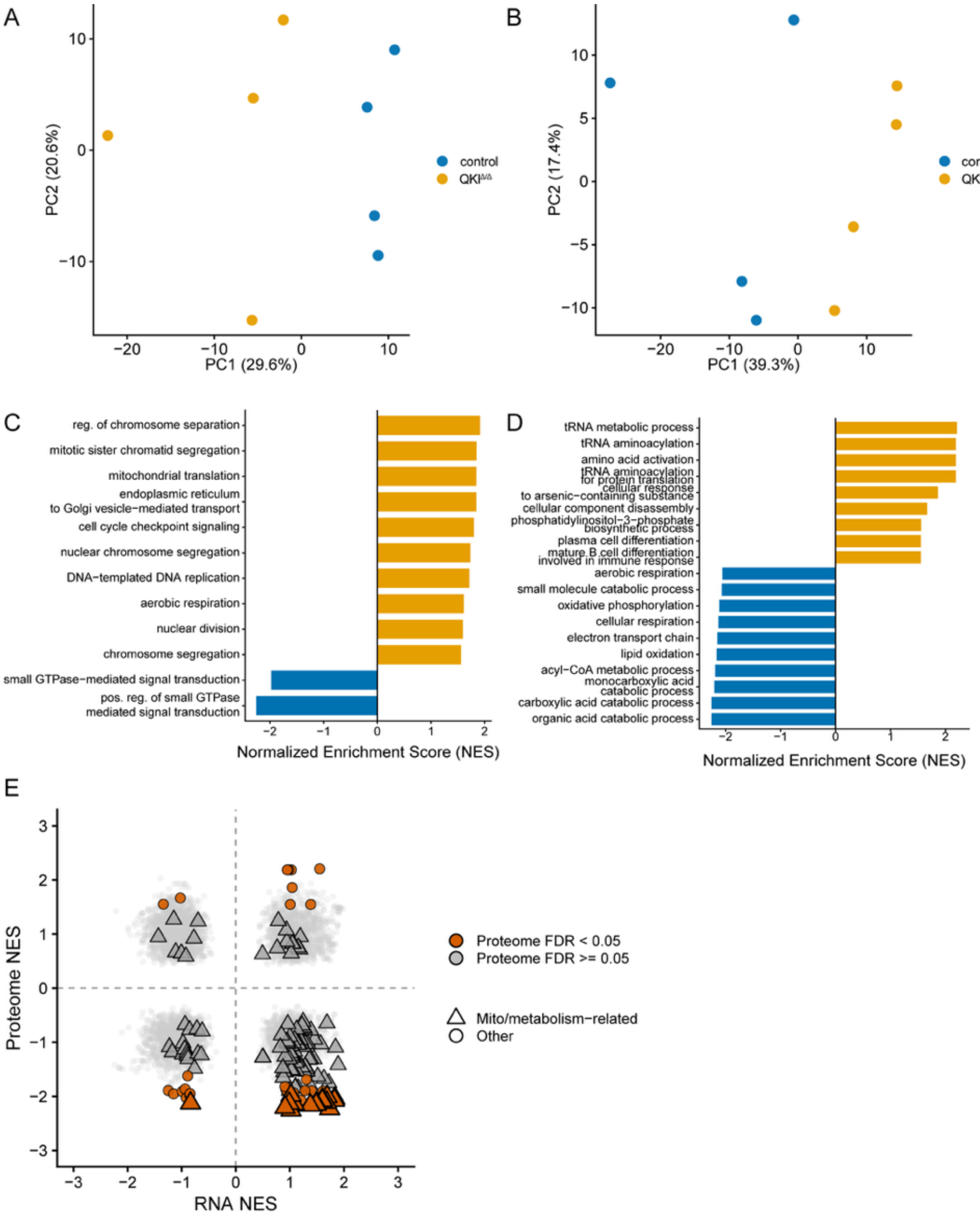


***Quality control and Gene Ontology Biological Process (GO-BP) sensitivity analyses for transcriptome–proteome integration***

**(A) RNA-seq principal component analysis (PCA) for quality control:** PCA of RNA-seq samples using variance-stabilized expression values from expressed genes. Points represent individual control and QKI$^{\Delta/\Delta}$ AT2 samples.

**(B) Proteome PCA for quality control:** PCA of proteomic samples using log2-transformed normalized protein intensities from protein groups quantified across all samples. Points represent individual and QKI$^{\Delta/\Delta}$ AT2 samples. A total of 2,043 protein groups were retained for PCA after excluding protein groups with zero or missing values in any sample and those with zero variance after log2 transformation.

**(C) RNA GO-BP gene set enrichment analysis (GSEA) for sensitivity analysis:** GO-BP GSEA of RNA-seq data using a protein-coding ranked gene list. Bars indicate NES for selected significant GO-BP terms after redundancy reduction using simplify.

**(D) Proteome GO-BP GSEA for sensitivity analysis:** GO-BP GSEA of proteomic data. Bars indicate NES for selected significant GO-BP terms after redundancy reduction using simplify, highlighting metabolic and respiratory processes.

**(E) RNA–proteome GO-BP normalized enrichment score (NES) scatter plot for sensitivity analysis:** Integrated comparison of RNA-seq and proteomic GO-BP GSEA results. Each point represents one GO-BP term matched by GO term ID. The x-axis shows RNA-seq NES, and the y-axis shows proteome NES. Terms significant in proteomic GO-BP

GSEA are highlighted, and mito/metabolism-related terms are indicated using keyword-based annotation.

***Figure E6***

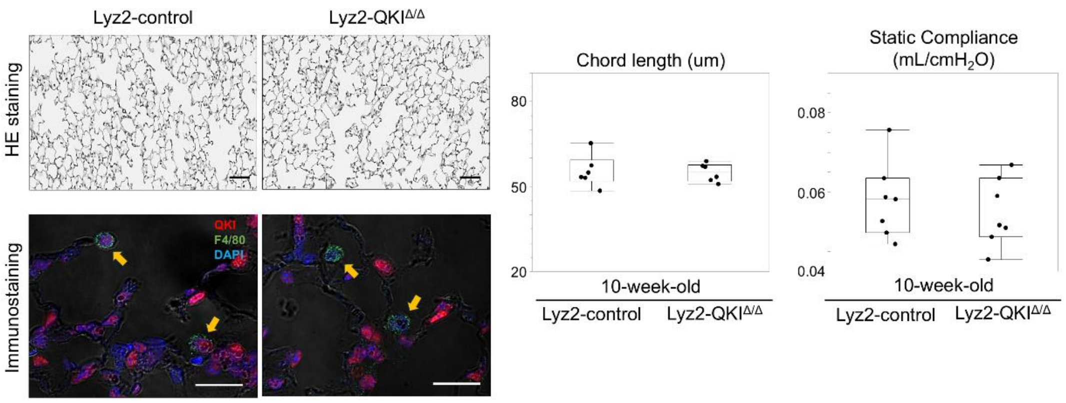


***Myeloid cell–specific QKI deletion does not increase airspace size or alter pulmonary mechanics in mice***

To evaluate the role of QKI in myeloid-lineage cells, including alveolar and interstitial macrophages, lung morphometry and pulmonary mechanics were analyzed in 10- to 12-week-old *Lyz2*$^{+/tm1(cre)Ifo}$ /*Qki*$^{flox/flox}$ (Lyz2- QKI$^{\Delta/\Delta}$) and *Qki*$^{flox/flox}$ (Lyz2-control) mice. In this model, QKI deletion was driven by Lyz2-Cre, resulting in predominant targeting of myeloid-lineage cells.

Representative microscopic images: Top, hematoxylin and eosin (H&E) staining of lung tissues. Scale bars, 50 μm. Bottom, immunofluorescence staining of lung sections for QKI (red), F4/80 (green; macrophage marker), and DAPI (blue). Yellow arrows indicate alveolar macrophages. Scale bars, 50 μm.

Quantification of alveolar chord length in 10-week-old control and Lyz2- QKI$^{\Delta/\Delta}$ mice was performed using a semi-automated method based on deep learning segmentation. n = 6 mice per group.

Pulmonary function parameters, including static compliance (Cst) were measured using a FlexiVent system in 10-week-old control and Lyz2- QKI$^{\Delta/\Delta}$ mice. n = 6 mice per group.

Box-and-whisker plots show individual data points. Statistical analyses were performed using Student’s t-test.

13. Haczku A, Atochina EN, Tomer Y, Cao Y, Campbell C, Scanlon ST, et al. The late asthmatic response is linked with increased surface tension and reduced surfactant

protein B in mice. *American Journal of Physiology-Lung Cellular and Molecular Physiology.* 2002;283(4):L755–L65.

30.
